# Brain-wide neural co-activations in resting human

**DOI:** 10.1101/2021.12.03.471116

**Authors:** Lei Ding, Guofa Shou, Yoon-Hee Cha, John A. Sweeney, Han Yuan

## Abstract

Spontaneous neural activity in human as assessed with resting-state functional magnetic resonance imaging (fMRI) exhibits brain-wide coordinated patterns in the frequency of <0.1Hz. However, fast brain-wide networks at the timescales of neuronal events (milliseconds to sub-seconds) and their spatial, spectral, and propagational characteristics remain unclear due to the temporal constraints of hemodynamic signals. With milli-second resolution and whole-head coverage, scalp-based electroencephalography (EEG) provides a unique window into brain-wide networks with neuronal-timescale dynamics, shedding light on the organizing principles of brain function. Using state-of-the-art signal processing techniques, we reconstructed cortical neural tomography from resting-state EEG and extracted component-based co-activation patterns (cCAPs). These cCAPs revealed brain-wide intrinsic networks and their dynamics, indicating the configuration/reconfiguration of resting human brains into recurring and propagating functional states, which are featured with the prominent spatial phenomena of global patterns and anti-state pairs of co-(de)activations. Rich oscillational structures across a wide frequency band (i.e., 0.6Hz, 5Hz, and 10Hz) were embedded in the dynamics of these functional states. We further identified a superstructure that regulated between-state propagations and governed a significant aspect of brain-wide network dynamics. These findings demonstrated how resting-state EEG data can be functionally decomposed using cCAPs to reveal rich structures of brain-wide human neural activations.

## Introduction

Spontaneous fluctuations are a hallmark of neural signals, which have been observed in electrophysiological^1,2^, hemodynamic^3,4^, and optical imaging^5^ studies in various species under different behavioral conditions^6,7^. These spontaneous fluctuations exhibit inter-regional functional connectivity at multiple spatial scales ranging from function-specific networks to the entire whole brain network^8^. Spatial and dynamic structures of these multi-level brain networks have been observed with abnormalities in almost all major neuropsychiatric disorders^9^, indicating their significant clinical value.

Brain-wide functional networks have been predominantly probed through noninvasive measurement of spontaneous hemodynamic signals^10^ using functional magnetic resonance imaging (fMRI)^11^. It has been assumed that inter-regional correlations observed in hemodynamic signals, reflecting spatial organizations of brain-wide intrinsic networks and their state-dependent changes, are converted from coordinated large-scale spatiotemporal dynamics of spontaneous neural activity via neurovascular coupling^12,13^. To date, multiple heterogeneous dynamics of large-scale spontaneous neural networks have been reported in animals using optical imaging^14,15^ and neurophysiological studies^16,17^ based on observations of propagation patterns from recordings of limited spatial coverage, i.e., restricted imaging areas^15,18^ and/or a few selected but largely-separated recording sites^19^. In human, a scalp-based electroencephalography (EEG) study^20^ has suggested a long-range anteroposterior propagation. Simultaneous EEG and fMRI recordings in human^21^ have demonstrated that regional EEG patterns are correlated with distributed hemodynamic activations^22^ and, interestingly, EEG signals from different scalp sites also show similar correlation patterns toward hemodynamic signals^18,23^. While indirect and limited, these phenomena attest to the existence of brain-wide neural networks in humans that underlie brain-wide hemodynamic networks. Unfortunately, there is no direct visualization of brain-wide neural networks in humans so far. More importantly, while brain networks may be best defined spatially by fMRI in terms of their anatomy, to investigate their temporal organization patterns, it requires high temporal resolution data such as EEG (e.g., 1000Hz) over fMRI (∼1 Hz).

The gap between brain-wide neural and hemodynamic networks is intrinsically tied to the difference between neuronal and vascular structures. It is largely unknown how observed global propagations among neurons, e.g., traveling wave^15,20^ and spiral wave^14^, are converted into the dynamics of vascular networks^24^. Motivated to understand this gap, a strategy shift of fMRI data analysis from traditional correlation-based approaches to ones integrating amplitudes has revealed several new brain-wide network patterns. This includes transient co-activation patterns (CAPs)^25-28^ among a subset of anatomically connected cortical areas, i.e., a prominent feature similarly observed in hemodynamic correlation structures, and brain-wide propagation patterns^28-30^, bridging more toward neuronal waves. A novel wide-field optical imaging study concurrently monitoring calcium signals reflecting neuronal spiking activity and hemodynamic signals in mice^29^ indicates that transient neural CAPs are embedded in brain-wide propagation patterns. This observation suggests that these two new brain-wide patterns are associated and brain-wide propagation patterns might play a critical role in formulating hemodynamic correlation structures.

Currently, optical imaging systems sensitive to neural activities cannot effectively penetrate human skulls^31^ preventing human studies from directly reporting and visualizing cortical-level brain-wide coordinated neural electrical phenomena and dynamics. The present study was conducted to identify such patterns on the human cortex from classical EEG data using an advanced computational framework developed based on state-of-the-art signal processing techniques. Our data indicate that brain-wide CAPs could be probed from high-density scalp-based EEG data and their cortical constructs could be directly visualized in the form of reconstructed tomographies. Our data further indicate rich dynamic structures in identified brain states at multiple time scales, including recurring, propagating, and oscillatory patterns. Finally, we report a superstructure involving all identified brain states that regulates between-state spatial, temporal, and transitional relationships and leads to characteristic propagation patterns coordinated by a pair of global co-activation brain states identified in all individuals.

## Methods

### Dataset and preprocessing

The experiment of the main dataset was approved by the Institutional Review Board at the University of Oklahoma Health Science Center (OUHSC), and written informed consents were obtained from all healthy participants. Resting-state EEG data (Dataset 0: 10 minutes long, n=34, 10 females, age: 24±5 years) with eye-closed were recorded at a sample frequency of 1000Hz using the 128-channel Amps 300 amplifier (Electrical Geodesics Inc., OR, USA). No sleep was noted as monitored by experimenters and/or reported by participants. Structural MRI was collected for each individual on a GE MR750 scanner at OUHSC MRI facility, using GE’s “BRAVO” sequence: FOV=240 mm, axial slices per slab = 180, slice thickness = 1 mm, image matrix = 256×256, TR/TE = 8.45/3.24ms. In addition, EEG sensor positions and three landmark fiducial locations (i.e., nasion, left and right pre-auricular points) were digitized by the Polhemus Patriot system. Two other datasets (see details in **Supplementary Note 1**) from healthy participants of our previous studies were included to validate the findings from Dataset 0. Briefly, dataset 1^60^ had eye-closed resting-state EEG data (5 minutes long, 128 channels, and sampled at 512Hz; n=19, 13 females, age: 13±6 years) and no individual structural MRI, where age-appropriate MRI templates^61^ were used. Dataset 2^62^ had eye-closed resting-state EEG data (5 minutes long, 126 channels, and sampled at 1000Hz; n=20, all females, age: 49±7 years) and individual structural MRI data. These three datasets exhibited a significant age difference across groups (*F*_(2)_ = 208, *p*<1e-6). All three EEG datasets were first filtered by a notch filter at 58 to 62Hz and a band-pass filter at 0.5–100Hz. Noisy channels interpolation and ICs removal related to ocular, muscular and cardiac activities were conducted using the EEGLAB toolbox^63^. Finally, EEG data were down-sampled to 250Hz and re-referenced to the common average. It is noted that no EEG segments were rejected to maintain the continuity of data for subsequent dynamic analysis.

### Cortical Source Imaging: Cortical Current Tomography

Cortical source imaging was performed individually to reconstruct cortical sources from scalp-level EEG signals (**Fig. 1A**). FreeSurfer^32^ was used to segment individual MRI data to extract the surfaces of the scalp, skull, and brain for volume conduction model, and the interface between white and gray matters for cortical current density (CCD) source model. The surfaces of volume conduction model were each tessellated into triangular elements of 10,242 nodes and 20,480 triangles, while the surface of the CCD model was tessellated into triangular elements of 20,484 nodes and 40,960 triangles. On the CCD model, the nodes on the medial wall adjoining the corpus callosum, basal forebrain, and hippocampus were excluded, and the total number of source nodes was reduced to 18,715. Each of these nodes was assigned with a dipole with its orientation set as the normalized vector sum of the normal directions of all triangles sharing the node. The electrical conductivities of the scalp, skull, and brain were assigned as 0.33/Ωm, 0.0165/Ωm, and 0.33/Ωm, respectively. EEG sensor locations were registered on the scalp surface by aligning three landmark fiducial points from both EEG and MRI recordings. Based on these models, the boundary element method^65^ was used to build the forward relationship: **Φ**(*t*)=**L**·**S**(*t*), where **L** is the lead field matrix; **Φ**(*t*) and **S**(*t*) are functions of time for scalp EEGs and dipole amplitudes, respectively. The minimum-norm estimate^66^ was used to reconstruct dipole amplitudes on the cortical surface: **S**(*t*)=**L**^T^·(**L**·**L**^T^+*λ*·**I**)^-1^·**Φ**(*t*), where *λ* was the regularization parameter and selected via the generalized cross validation method^67^ and **I** was the identity matrix. To control the quality of reconstructed cortical sources, the selected *λ* values beyond three standard deviations of all values of each individual were considered as outliers and interpolated with the neighboring ones. Based on these adjusted *λ* values, cortical current tomography was reconstructed as a function of time for each participant for subsequent analysis.

**Figure 1.**
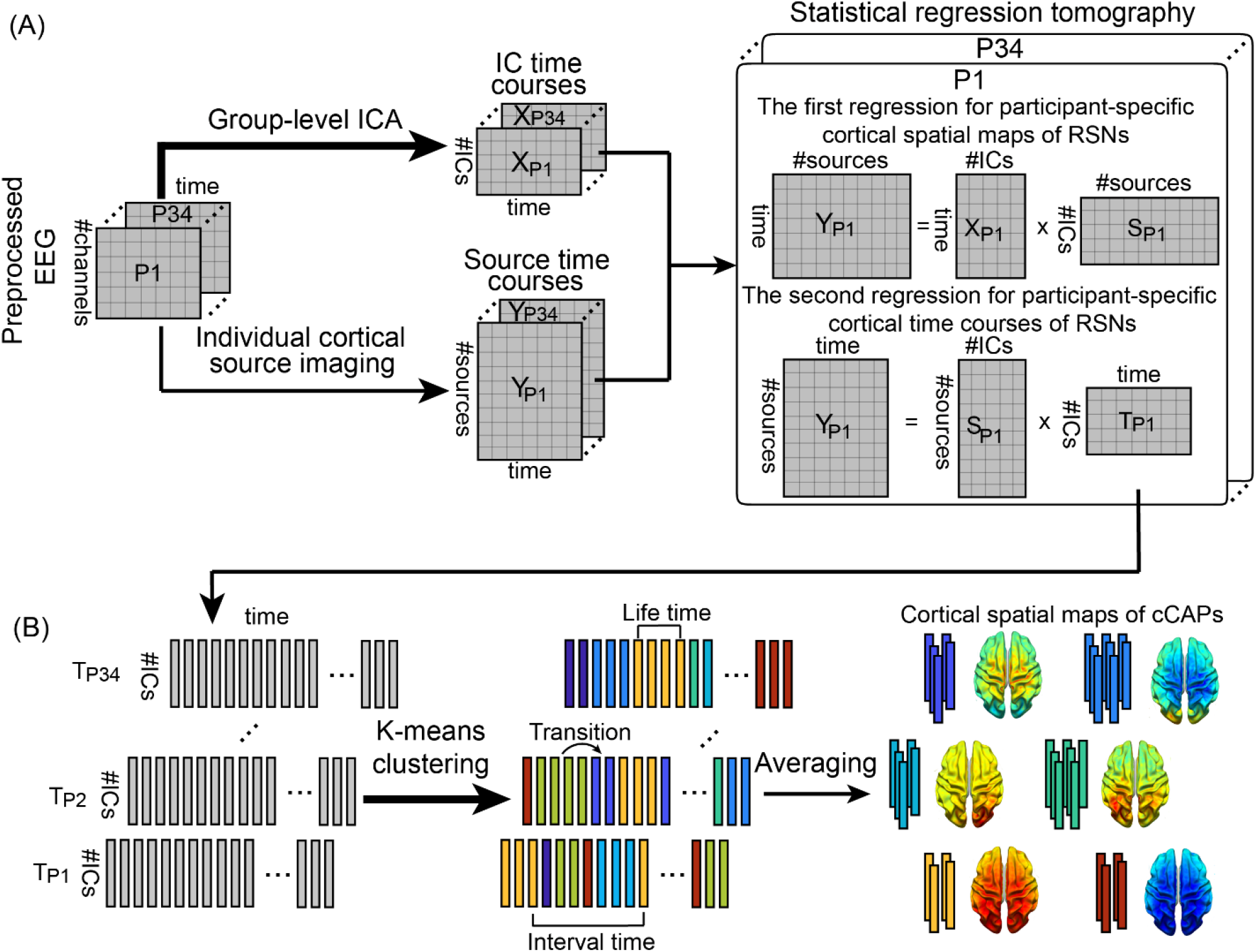
Schematic illustration of the method. (A) Spatial definitions of cortical RSNs and their dynamics calculated using a statistical dual-regression analysis between sensor-level IC time courses and cortical-level source time courses. IC time courses are calculated from a group-level ICA on preprocessed EEG data temporally concatenated across all participants. Cortical source time courses are calculated by cortical source imaging from EEG and MRI data of individual participants. (B) Recurring brain states and their dynamic transitions can be captured and classified by clustering timeframe data of cortical RSNs into short-lived spatially congruent patterns (i.e., cCAPs) using the k-means algorithm. P: participant; T: timecourse.

### Group-level ICA: scalp-level representations of RSNs

A group-level ICA was used to decompose preprocessed scalp EEG data into multiple ICs^60,62^. Briefly, individual EEG data were normalized to z values and temporally concatenated. A short-time Fourier transform was then performed on segmented 1-s epoch data without overlap to obtain time-frequency representations of EEG data on channels, which were the input for the time-frequency ICA model^68,69^. The Fourier transform modulated rhythmic neural activities that were usually Gaussian distributed into more super-Gaussian to be better detected from artifacts^68,69^. Here, the Fourier spectrum data were selected in the range of 1 to 100Hz at a resolution of 1Hz, which were individually normalized to account for the 1/f distribution over the spectrum. The ICA model was run for each model order from 25 to 64 with 64 finally being selected as the order of the ICA model for subsequent analyses, as it gave the best spatial patterns in identifying major RSNs. To obtain IC time courses, original EEG data were projected using the demixing matrix calculated from the group-level ICA. Thereafter, ICs showing neural activation characteristics in both of their spatial and spectral patterns, as compared to the ones reported in the literature^22,33,69^, were selected as the scalp-level representations of cortical RSNs.

### Statistical Regression Tomography: cortical representations of RSNs

For each participant, the cortical representation of each RSN was estimated via a statistical dual-regression analysis between time courses of individual cortical dipoles and the time courses of the selected ICs (**Fig. 1A**). First, both time courses were down-sampled to 100Hz and their envelope data were calculated via the Hilbert transform^2^. Second, to obtain cortical maps of RSNs, the first regression was performed with time courses of all selected ICs as the regressors and time courses of individual dipoles as the response data. In this step, an autoregression (AR) model with the order of 6 was used to remove the autocorrelation in the regressors^70^. Third, the second regression was performed to reconstruct time courses of cortical RSNs with cortical RSN maps obtained from the first regression analysis as the regressors and timeframe-wise spatial maps of cortical dipoles as the response data. As a result, cortical tomographies of RSNs were defined with their corresponding spatial and temporal patterns in individuals.

### Identifying component-based CAPs: A K-means clustering analysis

Time courses of cortical RSNs obtained above were subject to a K-means clustering analysis to identify distinct patterns of their temporal co-activations (**Fig. 1B**), which was termed as component-based CAPs (cCAPs) as these time courses represented dynamic fluctuations of IC signals from EEG. The clustering analysis was performed at the group level on matrix data, i.e., number of time points from all participants × number of RSNs. To account for different variances in different RSNs and participants, each time course was normalized as zero-mean and unit-variances before the clustering analysis. In the clustering model, the L1-norm distance was used as the metric to measure the similarity among timeframe-wise data and, therefore, the co-activation here was defined on amplitudes, rather than temporal correlations, similar to other CAP studies in fMRI^25,27,28^. After calculating the model order in a range from 2 to 20 for clustering, we chose the model order of 8 to report our results as it was a good tradeoff between observing more distinguishable cCAPs and producing more similar cCAPs. The output of the clustering analysis labeled each timeframe data of cortical RSNs to a unique cCAP (**Fig. 1B**). Thereafter, the cluster centers of individual cCAPs were obtained as the vectors of weights of selected RSNs via averaging original values of cortical RSN timeframe data (before normalization) with the same corresponding labels (**Fig. 1B**). The weights in these center vectors were further numerically ranked, i.e., from 1 to 8 in their absolute values, for each RSN across all eight cCAPs to illustrate relative activation levels of each RSN among all cCAPs. The distances between these center vectors were calculated and projected into a 3D space (**Fig. 2C**) using a multidimensional scaling tool from MATLAB (i.e., cmdscale.m) to examine the spatial relationship among all cCAPs. Finally, the cortical tomography of each cCAP was built for individuals as the weighted sum of cortical tomographies of RSNs from individuals by the corresponding center vector (**Fig. 1B**). Using the same means, cortical tomographies of all cCAPs at each timeframe were similarly reconstructed to show their activation sequences in individuals as in **Fig. 4** and **Supplementary Movies 1-4**.

### Temporal metrics of cCAPs

Multiple temporal metrics (**Fig. 1B**) were calculated on data from individuals and then summarized to generate group-level statistics, e.g., means, standard deviations, and histograms. The metric of *lifetime* was defined as the duration of each occurrence of a cCAP. A *transition* happened when two neighboring timeframes were labeled with different cCAPs. The *interval time* of a cCAP was defined as the time difference between its two consecutive occurrences from the end of the early one to the beginning of the late one. To probe temporal long-scale relationship between the occurrences among all eight cCAPs, an alignment analysis was developed, where one cCAP was selected as a reference cCAP and then the occurrence probabilities of all cCAPs at certain distances in the time axis were calculated with respect to the reference cCAP. Specifically, the epochs of all occurrences of the reference cCAP were extracted each with 5 seconds before the start of and 5 seconds after the end of an occurrence. Within the total 10 seconds for each epoch, the occurrences of all eight cCAPs at each time instant were calculated in individuals and then divided by the total number of epochs to obtain their occurring probabilities, each presented as a function of time centered toward the reference cCAP.

### Propagation patterns among cCAPs

To investigate the propagation structures among all cCAPs, two analyses were performed on individual’s data. First, one-step transition probabilities of a cCAP to other cCAPs were defined as the numbers of transitions from the cCAP to other cCAPs (i.e., outflow) or from other cCAPs to the cCAP (i.e., inflow), divided by the total number of occurrences of the cCAP. These probability data were used to form the outflow/inflow matrices with the current cCAP state at time *t* in the vertical direction and the next cCAP state at time *t*+1 in the horizontal direction, in which the elements in any row of the outflow matrices summed to one and the elements in any column of the inflow matrices summed to one, known as percentage outflow/inflow rates of cCAPs. Second, we studied the propagations among distinct functional brain states (i.e., cCAPs) coordinated by two polarized cCAPs, i.e., cCAPs 7 and 8 (see Fig. 2C and Results). For the purpose of comparisons, two contrasting propagations between cCAPs 7 and 8 themselves were also defined. Therefore, the entire recording for each participant was segmented into four types of propagation intervals: cCAP 7→8, cCAP 8→7, cCAP 7→7, and cCAP 8→8. For these four types of propagations, multiple metrics were calculated based on data from individual propagations in each participant and statistically compared using repeated measures ANOVA (rmANOVA) and *t*-test when appliable (Fig. 6), including occurrences (Fig. 6A), means and histograms of propagation durations (Fig. 6B), number of cCAPs visited (counting repeated visits to the same cCAP) and number of different cCAPs visited (other than cCAPs 7 and 8) in each propagation (Fig. 6C), and occurrence rate histograms of four propagation types as the function of number of different cCAPs visited (Fig. 6C). To study the roles of the rest six cCAPs in these four propagation types, we calculated the occurrence rates of propagations with the visit to a specific cCAP out of total propagations of the same type for all four types of propagations, separately (Fig. 6D). We further broke down these occurrence rate data according to the number of different cCAPs visited (Fig. 6E). To investigate whether the propagation patterns coordinated by cCAPs 7 and 8 were unique, same types of propagations based on two *pseudo*-polarized states made by all other possible pairs (total 27 pairs) of the eight cCAPs were examined and all above-mentioned calculations were repeated. Unless specified, *p* values were corrected by Bonferroni method.

## Results

### Brain-wide EEG Component Clustering Analysis Reveals a Set of Spatially-structured Functional Brain States

Using concatenated high-density EEG resting-state data from 34 participants (128 channels, sampled at 1000Hz, Dataset 0), mapped onto participant-specific models of cortical surfaces from FREESURFER^32^ by our newly developed computational framework (**Fig. 1**), we identified eight reproducible CAP patterns from EEG component signals (component-based CAPs, cCAPs, **Fig. 2A**), where these component signals have been related to the definitions of electrophysiological resting state networks (RSNs, **Supplementary Fig. 1A**) in previous studies^33,34^. Essentially, this framework finds, in a completely data-driven way, recurrent states of networked activities (i.e., cCAPs) in human brains with structured spatiotemporal properties. Spatially, almost all cCAPs (with the exception of cCAP 3 relatively focusing on the cingulate cortex) suggest brain-wide patterns of co-(de)activations in both their cortical maps (**Fig. 2A**) and activity levels of involved RSNs (**Fig. 2B** and **Supplementary Note 2**). These facts indicate that identified cCAPs activate anatomically-connected and functionally-related neural substrates in the dynamic behaviors of the resting human brain. Two cCAPs (cCAPs 7 and 8) indicate global co-activation and co-deactivation patterns, respectively, as all RSNs reach their own top (or close to top) levels of positive or negative magnitude of activity, which are further supported by their cortical maps. Moreover, default mode networks (DMNs, **Supplementary Fig. 1A**) and task-positive networks (TPNs, RSNs other than DMNs) reveal opposite co-activation patterns where high-magnitude DMN activations are accompanied by relatively low-magnitude TPN activations in cCAPs 2, 5 and 6 (**Fig. 2B**) and vice versa in cCAPs 1, 3, and 4. As a notable feature of CAPs identified in fMRI, the configuration into anti-state pairs is characterized by opposing patterns of functional co-(de)activations^28^. We conducted a sequential search for such pairs based on the metric of vectorized spatial correlation coefficients. Apart from the hemisphere-mirrored pair (cCAP 2-6), the anti-state characteristic is especially prominent in the cCAP 7-8 pair (*r* = - 0.90±0.05), but also apparent in the cCAP 1-4 pair (*r* = -0.41±0.23) and the cCAP 3-5 pair (*r* = - 0.27±0.18). We further calculated spatial distances among vectorized cortical patterns of all eight cCAPs and projected these vectors into a 3D space (see Methods). The results reveal a well-structured spatial relationship among eight cCAPs (**Fig. 2C**), in which cCAPs 7 and 8 are positioned as two poles with the longest distance and the other six cCAPs are clustered halfway between them. Furthermore, cCAP 5 is significantly closer to cCAP 8 while both cCAPs 2 and 6 are significantly closer to cCAP 7 than other cCAPs (**Supplementary Note 2**). The vectorized spatial correlation coefficients of eight cCAPs between those identified in individuals and the group-level ones indicate that these cCAPs can be reliably detected in individuals (**Supplementary Fig. 2C**). In particular, cCAPs 7 and 8 have the highest spatial correlations (only two *r* > 0.5) (**Supplementary Fig. 2B**).

**Figure 2.**
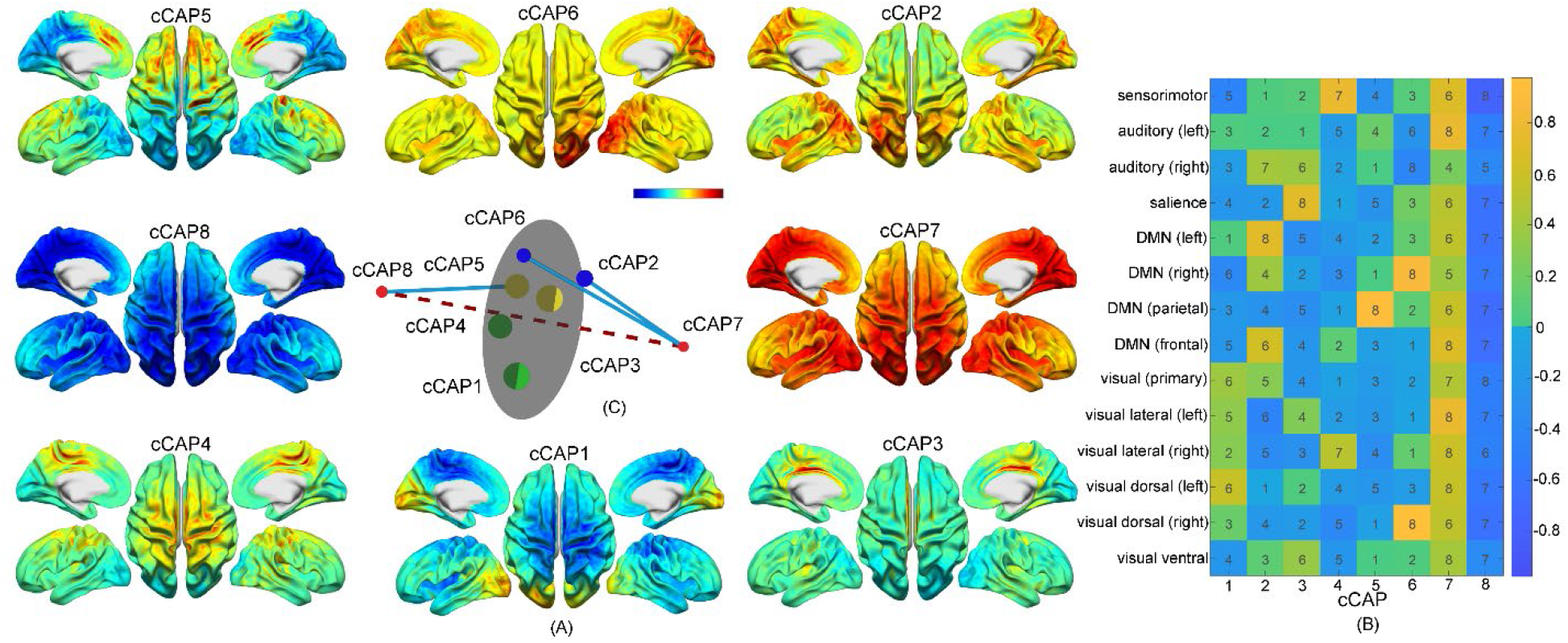
A set of spatially-structured functional states with brain-wide patterns, i.e., cCAPs, from the resting human brain. (A) Cortical maps of the cCAPs identified at the group level in which both cCAPs 7 and 8 show global co-(de)activation patterns. Red-yellow colors indicate co-activations (i.e., high neuronal currents), and blue indicates co-deactivations (i.e., low neuronal currents). (B) The weight vectors, i.e., columns, of the cluster centers of all cCAPs. Colors indicate weight amplitudes (with ± signs) of individual RSNs at the cluster centers of cCAPs and numbers indicates the amplitude ranks (no signs) of same individual RSNs across all cCAPs. DMN: Default mode network. (C) The distance map of the weight vectors of all cCAPs projected into a 3D space. Same-color dots: anti-state pairs and hemisphere-symmetric pair; red dots: two polarized states, i.e., cCAPs 7 and 8, connected by the dashed line; blue lines: connecting brain states that are structurally closest to two polarized states (see more in **Supplementary Fig. 2A**); gray circular plane: halfway between two polarized states.

### Recurring Patterns of cCAPs Support the Spatial Structure Formulated by the Set of Brain States

We then investigated cCAP dynamics with the goal of finding evidence to support the observation of multi-level spatial structures formulated by the eight cCAPs. We report the occurrence rates of individual cCAPs that are within a range of 9% and 15% (**Fig. 3A**), close to the equal opportunity of occurrence (i.e., 12.5% for eight cCAPs). The lifetime of each cCAP occurrence was in the range between 25 and 35 milliseconds for mean values (**Fig. 3C**). Both data suggest all cCAPs are recurring states that can be reliably detected at both group and individual levels as their variations among individuals are low comparing with their group-level mean values (**Figs. 3A**, **3C**), which are consistent with individually reproducible cCAP spatial patterns discussed above. The cCAPs 7-8 pair reveals significantly lower occurrence rate (*p<0.0005, FDR corrected*), longer lifetime (*p<0.00005, FDR corrected*), and longer interval time between successive occurrences (*p<0.005, FDR corrected*) than all other cCAPs. The two anti-state pairs (cCAP 1-4 and cCAP 3-5) that are spatially close (**Fig. 2C**) share similar data for these temporal metrics. The hemisphere-mirrored cCAP pair (cCAP 2-6) have occurrence rates higher than the two polarized states (*p<0.005, FDR corrected*) but lower than the four anti-states (*p<0.0005, FDR corrected*). Their interval times are lower than the two polarized states (*p<0.05, FDR corrected*) but higher than the four anti-states (*p<0.0005, FDR corrected*). The lowest occurrence rates and longest interval times of cCAPs 7 and 8 are consistent with these two states being at the boundaries of distance maps (**Fig. 2C**), exhibiting low chances to be visited during between-state transitions (see **Figs. 4 and 5** below). Other states are more likely to be visited since they are closer to each other. In summary, the consistencies observed on cCAP spatial and temporal patterns at three levels, i.e., individual cCAPs, cCAP pairs, and the cluster of all eight cCAPs, suggest that these observations are manifested from the same underlying source, i.e., dynamically coordinated and networked brain-wide activations.

**Figure 3.**
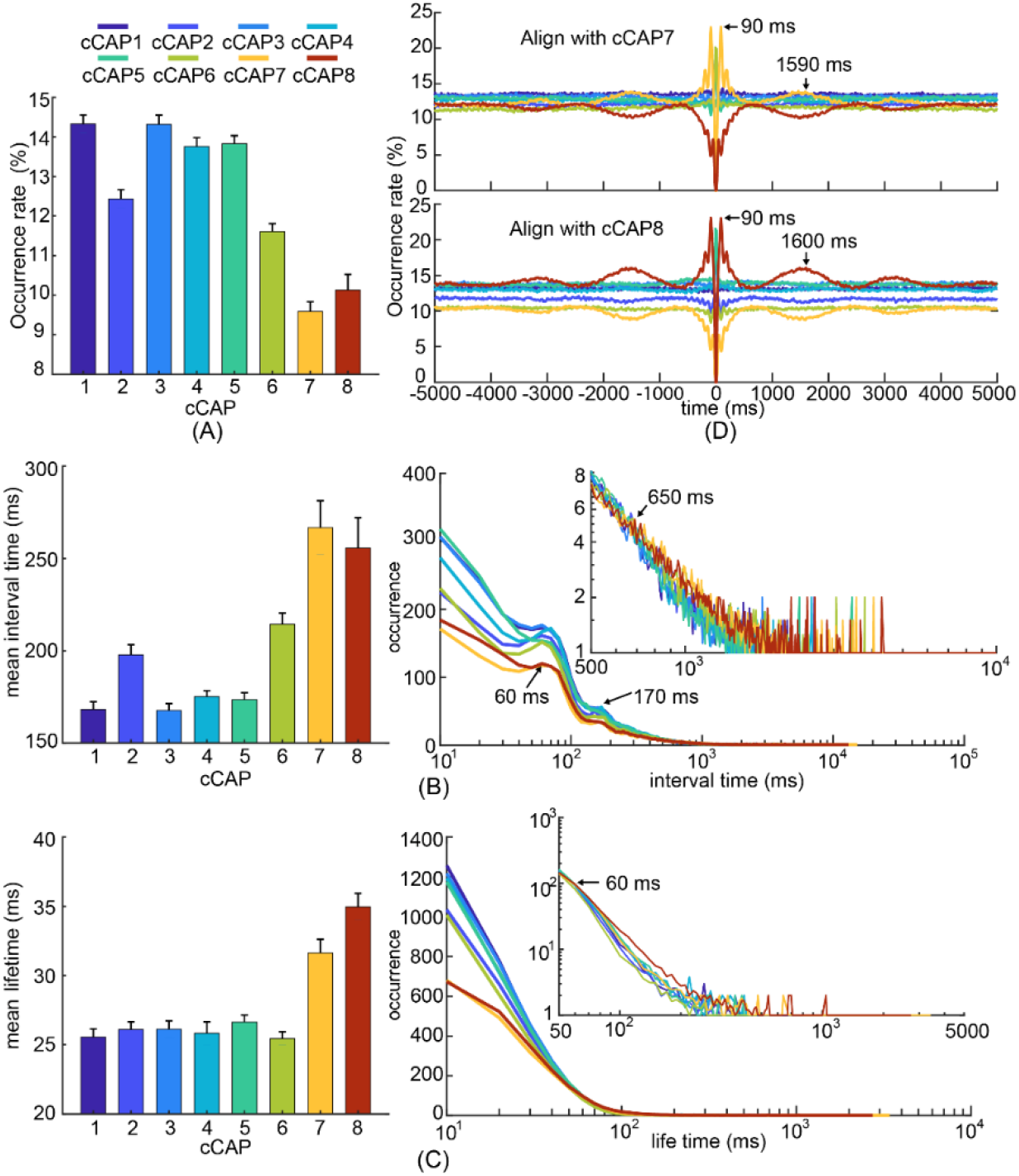
Temporal characteristics of recurring functional brain states, i.e., cCAPs. (A) Occurrence rates of cCAPs (mean±SEM) where cCAPs 7 and 8 show significantly lower occurrences than others (*p*<0.05, FDR corrected). (B) Mean (±SEM) values (left) and histograms (right) of interval times of cCAPs. Note that cCAPs 7 and 8 have significantly longer mean interval time than others (*p*<0.05, FDR corrected). All histograms show two peaks (around 60ms and 170ms) and the occurrences of cCAPs 7 and 8 become higher than other cCAPs beyond the interval time of 650ms (the inset). (C) Mean (±SEM) values (left) and histograms (right) of lifetimes of cCAPs. Note that cCAPs 7 and 8 have significantly longer mean lifetime than others (*p*<0.05, FDR corrected). The occurrences of cCAPs 7 and 8 become higher than other cCAPs beyond the lifetime of 60ms (the inset). (D) Occurrence probabilities of all cCAPs as functions of time distances toward all occurrences of the target cCAP 7 (top) and cCAP 8 (bottom). See other cCAPs as the target cCAP in **Supplementary** **Figs. 4B-C**. SEM: standard error of mean.

### Individual cCAPs Show Intrinsic Temporal Dynamics and cCAPs 7-8 Further Indicate Coordinated Large-Scale Fluctuations

The histograms of cCAP interval times (**Fig. 3B**) exhibit two characteristic peaks (∼10Hz and ∼5Hz, **Supplementary Note 2**) on top of exponentially decreasing curves. The wide ranges of interval times illustrated in the decreasing curves indicate that the occurrences of these recurring brain states are nonstationary, while two peaks reveal weak but observable intrinsic dynamic rhythms in cCAPs. These are consistently detected in individuals (**Supplementary Fig. 3A**). Note that the significantly longer interval times of cCAP 7-8 are achieved via having fewer short intervals and more long intervals (the inset, **Fig. 3B**) without lengthening the characteristic peaks. The same mechanism is also observed in cCAP 2-6 for their moderately but statistically significantly longer interval times (*p < 0.0005, FDR corrected*), as well as for longer lifetimes of cCAPs 7, 8, 2, and 6 (**Fig. 3C**). These observations suggest that both lifetimes and interval times of cCAPs are potentially modulated by some unrevealed mechanisms with longer temporal scales^28^ . To this end, we next examined long-scale temporal dynamics beyond interval times using an alignment analysis on occurrences of a target cCAP at different time distances toward all occurrences of a reference cCAP (see Methods). We observe two oscillatory phenomena elevated from the baseline of 12.5% (i.e., equal opportunity for eight cCAPs) that are exponentially decayed over the distance to the reference cCAP (**Fig. 3D** and **Supplementary Fig. 4A**). The short-scale oscillations have an inter-peak distance of ∼100ms, corresponding to the 10Hz frequency component in **Fig. 3B**. The long-scale oscillations show an inter-peak distance of ∼1.6 seconds, corresponding to a frequency of <1Hz. These elevated oscillatory phenomena could be reliably detected at both the group and individual levels (**Supplementary Fig. 4B**) but mainly when the target and reference cCAPs are same. One notable exception is the cCAPs 7-8 pair, in which elevated oscillations in one of them lead to symmetric but depressed oscillations in the other. This coordination between cCAPs 7 and 8, together with other data distinguishing them from the other cCAPs (**Figs. 2** and **3A-C**), suggests their important roles in modulating temporal dynamics of the resting human brain activity encoded in all cCAPs of brain-wide spatial patterns.

### Transition Patterns across cCAPs Support Spatial Structures Formulated by cCAPs

We then moved on to study between-state transitions via firstly investigating one-step transitions among cCAPs. The outflow matrix indicates patterns of lower transitions from all cCAPs to cCAPs 7 and 8 (last two columns, **Fig. 4A**) than to the other 6 cCAPs. Similarly, the inflow matrix indicates lower transitions from cCAPs 7 and 8 to all other cCAPs (last two rows, **Fig. 4B**). More prominently, the immediate transitions between cCAP 7 and 8 for both outflow and inflow are almost zero (<0.5%). All these observations support cCAP 7/8 as a polarized state similarly suggested in their distance maps and recurring patterns (**Figs. 2-3**). To find which one(s) from other six cCAPs have more immediate transitions to cCAP 7/8, percentage outflow rates (i.e., elements of a column, **Fig. 4A**) and percentage inflow rates (i.e., elements of a row, **Fig. 4B**) for each of these six cCAPs were compared, respectively, as actual outflow/inflow data biased by their occurrence rates (**Fig. 3**). cCAPs 2 and 6 show the maximal and significantly higher (as compared with the corresponding second largest) outflow rates (9% and 23% larger, respectively, both *p<0.05, corrected*) and inflow rates (14%, *p<0.05*, and 41% larger, *p<0.05, corrected*, respectively) toward cCAP 7. cCAP 5 shows the maximal and significantly higher outflow rates (15% larger, *p<0.05, corrected*) and inflow rates (24% larger, *p<0.05, corrected*) toward cCAP 8. All other three cCAPs (i.e., 1, 3, and 4) show no significant differences between the largest and second largest outflow/inflow rates. These observations are consistent with the distance map (**Supplementary Note 2**) where cCAPs 2/6 are closest to cCAP 7 and cCAP 5 is closest to cCAP 8.

**Figure 4.**
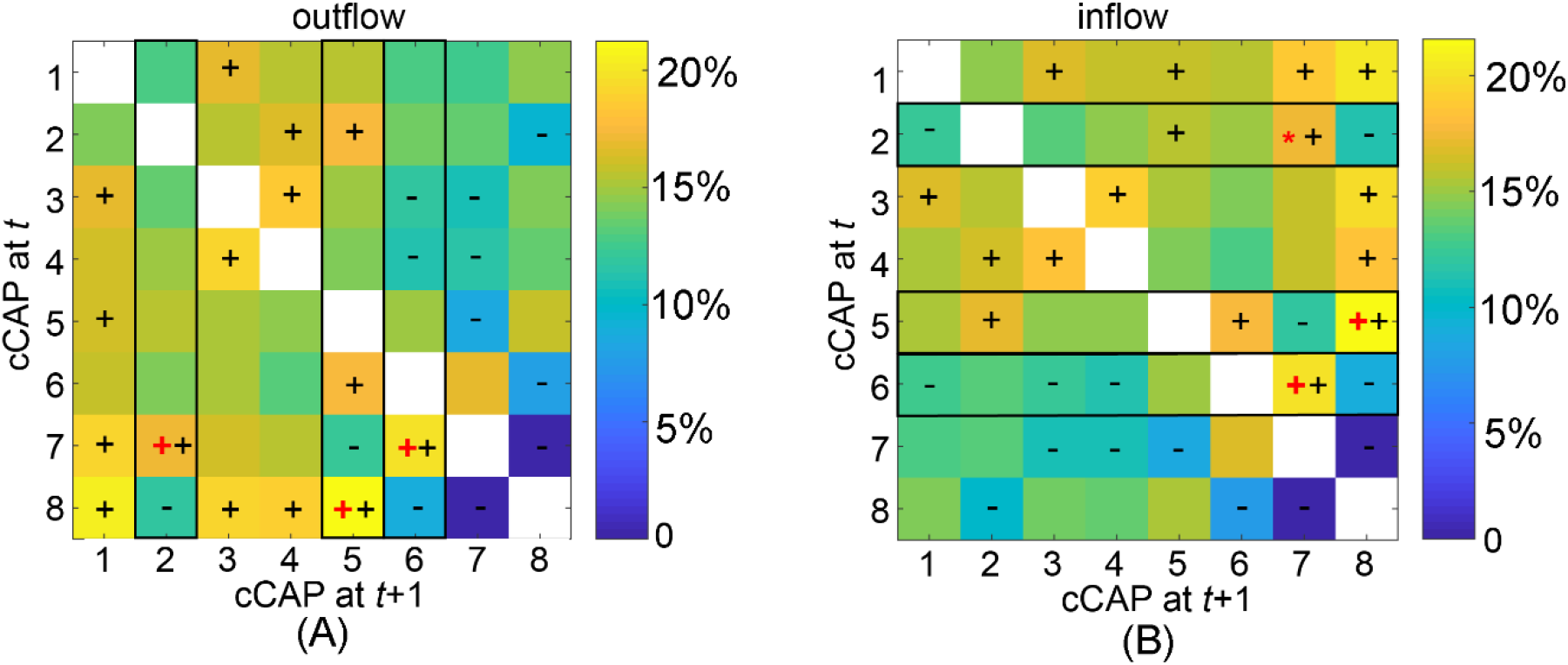
The immediate transition probability (one-step from time *t* to time *t*+1) among functional brain states: (A) the outflow matrix (normalized in rows) and (B) the inflow matrix (normalized in columns). The signs ‘+’ and ‘-’ denote significantly higher or lower transition probability values, respectively, than the equal probability value (i.e., 1/7, *p*<0.05, Bonferroni adjusted). The black rectangles highlight the columns (in A) and rows (in B), in which the cCAPs with the largest outflow or inflow values that are significantly higher than the second largest values are labeled (‘+’: *p*<0.001, Bonferroni adjusted, ‘*’: *p*<0.05, unadjusted). These identified cCAPs (labeled as ‘+’ or ‘*’) appear only to be the ones that are spatially close to two polarized states (Fig. 2C).

### Spontaneous Waves of cCAPs Show Unique Characteristic Patterns Frequently Propagated between Global Co-activation and Co-deactivation States

One goal of the present study was to understand how large-scale dynamics of neural activity is reflected in the spatial structure of cCAPs. When we visually inspected movies of spontaneous cCAPs (**Supplementary Movies 1-4** in continuous timeframes), we immediately noticed the frequent occurrence of waves of cCAPs that propagated between two polarized states (**Fig. 5**), either originating from cCAP 7 to cCAP 8 (cCAP 7→8) or vice versa (cCAP 8→7). These propagations occupy on average 41.1% (std: 3.0%) of total recordings from individual participants, with equal fractional occupation rates for both directions (cCAP 7→8: 20.3%±1.7%; cCAP 8→7: 20.8%±1.6%). To gain insights about the propagations between cCAPs 7 and 8, the entire time courses of all cCAPs from individuals (of same lengths as original EEG recordings) were segmented into four types of propagation intervals: two propagations of interests: cCAP 7→8, cCAP 8→7 and two contrasting propagations: cCAP 7→7, cCAP 8→8 (see **Methods**). To answer whether the phenomena observed were only orchestrated by cCAPs 7 and 8, same types of intervals based on two *pseudo*-polarized states made by all other possible pairs (total 27 pairs) of the eight cCAPs were examined. We observe that the propagations between cCAPs 7 and 8 occur significantly lower (>85% lower, *p<1e-6, corrected*) (**Fig. 6A**), take about 1.6 times longer (*p<0.0005, corrected*, inset, **Fig. 6B**), and visit over 1.2 times more states (*p<1e-4, corrected*, inset, **Fig. 6C**) than the contrasting propagations (see more in **Supplementary Note 3**). The breakdown data according to the propagation duration (**Fig. 6B**) showed exponential decreasing patterns for four propagations and more occurrences of cCAP 8→7 and 7→8 for the durations over 100 ms (*post-hoc t* tests: *p<0.01 and consecutive durations>5*), and less occurrences for the durations below 100 ms than the contrasting propagations (*post-hoc t* tests: *p<0.01 and consecutive durations>5)*. A characteristic peak for both cCAP 8→7 and 7→8 appears around this point of separation (i.e., 100 ms) indicating the shift toward longer propagation times needed to visit more states (inset, **Fig. 6C**). The extremely low short-duration propagations (i.e., 10ms and 20ms) in cCAP 8→7 and 7→8 is consistent with the largest distance (**Fig. 2**) and almost zero one-step transitions (**Fig. 4**) between cCAPs 7 and 8.

**Figure 5.**
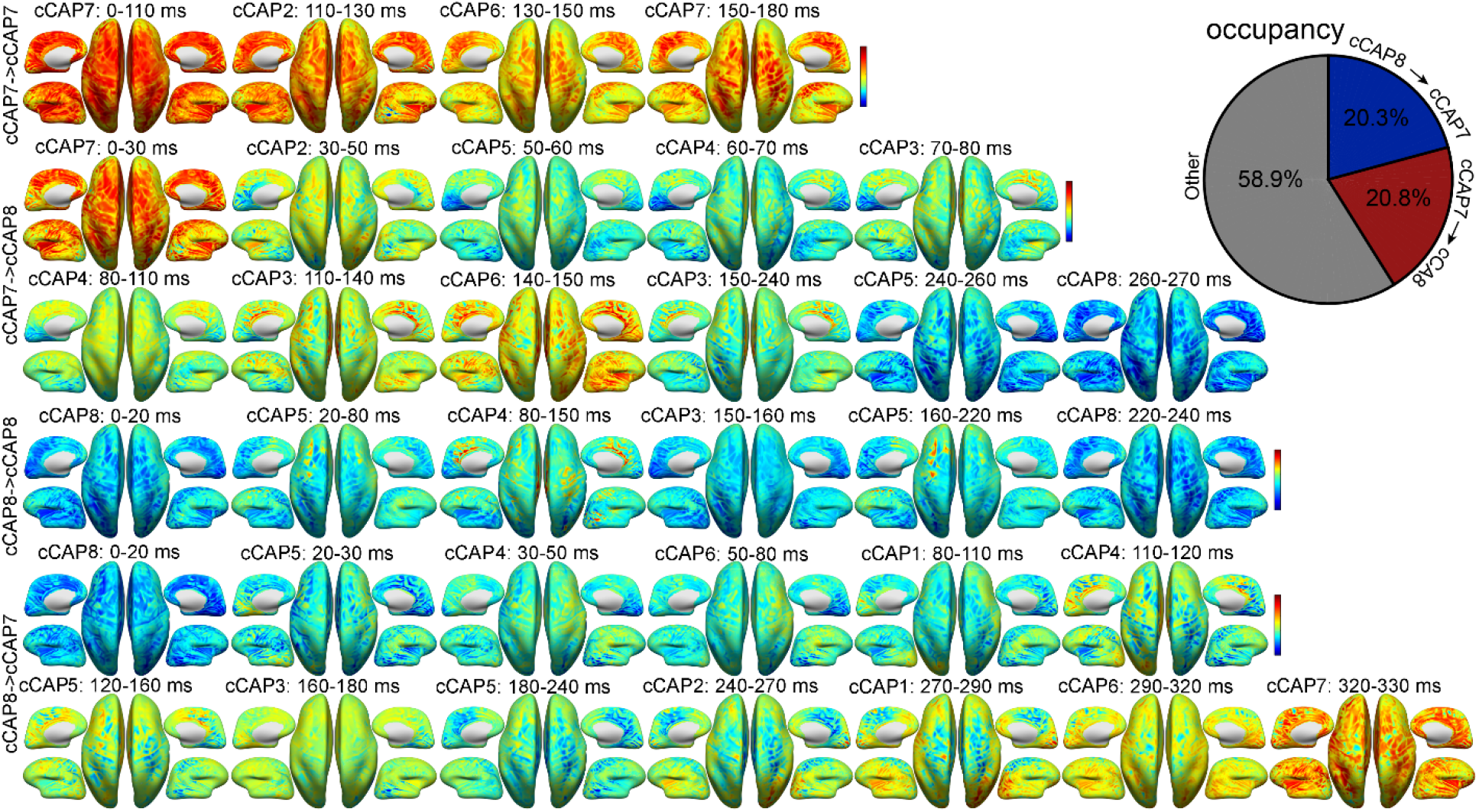
Examples of propagations between two polarized functional brain states (Fig. 2) on the inflated cortical surface from a participant: cCAP 7→7, cCAP 7→8, cCAP 8→8 and cCAP 8→7. Each map represents the averaged cortical pattern of an occurred cCAP over all timeframes within its lifetime window (labeled above) during the sequenced propagation. See Supplementary Movies 1-4 for corresponding videos in continuous timeframes. The pie plot indicates the total occupancies of the propagations between two polarized brain states, i.e., cCAP 7→8 and cCAP 8→7, over the entire recordings.

**Figure 6.**
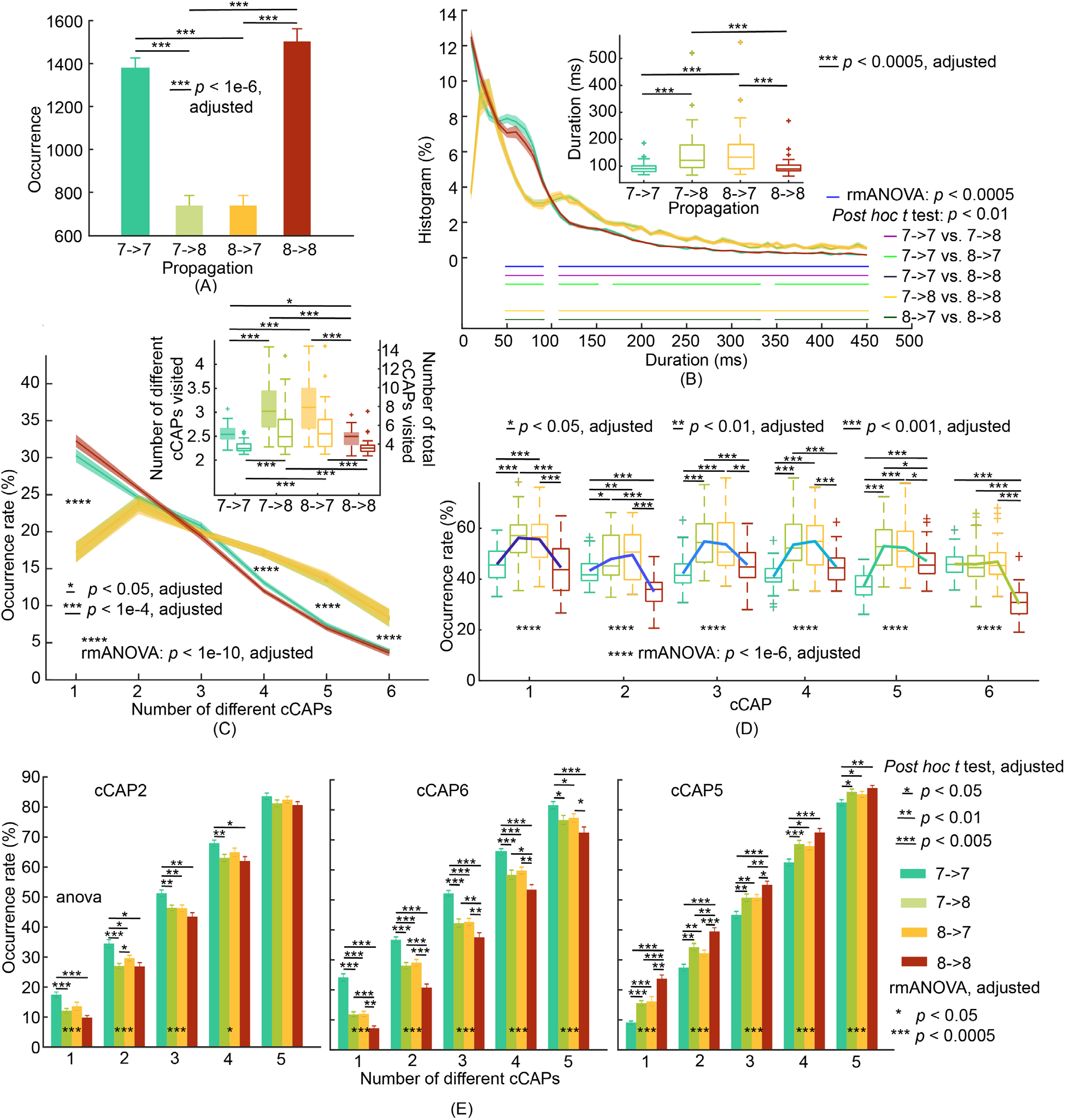
Patterns of propagations between two polarized functional brain states of cCAPs 7 and 8. (A) Occurrences of two propagations: cCAP 7→8 and cCAP 8→7, and two contrasting propagations: cCAP 7→7 and cCAP 8→8. (B) Boxplots (inset) of mean durations and histograms of durations (SEM: shaded areas) as functions of time for four propagations. (C) Occurrence rates of propagations (SEM: shaded areas) as functions of the number of different cCAPs (the six cCAPs other than cCAPs 7 and 8) visited per propagation. Inset: Participant-level means (±SEM) of numbers of different cCAPs (left y-axis and boxplots with solid-fills) and total numbers of cCAPs (right y-axis and boxplots with no-fills) visited per propagation. Post-hoc *t* tests indicate significant differences (*p*<0.01, adjusted) when the number of different cCAPs equals to 1, 4, 5, and 6 between the propagations of cCAP 7→8 and cCAP 8→7 and the two contrasting propagations. (D) Occurrence rates of the other six cCAPs within each type of propagation. (E) Occurrence rates of cCAPs 2, 6 and 5 within each type of propagation as functions of number of different cCAPs visited per propagation. See **Supplementary** Fig. 6 for cCAPs 1, 3, and 4. The condition of number of cCAPs as 6 is omitted since all occurrence rates are 100% by the definition of this metric.

It is further suggested that the propagations between cCAPs 7 and 8 not only visit more other states, but also visit more different states (**Fig. 6C**). Percentage-wise, significantly more cCAP 8→7 and 7→8 than the contrasting propagations (rmANOVA: *p<1e-10, corrected; post-hoc t tests: p < 0.001, corrected*) happen when the numbers of different states visited are high (i.e., 4, 5, and 6). When only one other state is visited, significantly less cCAP 8→7 and 7→8 (*p<1e-10, corrected*) happen than the contrasting propagations. We further studied the occurrence rates of other six cCAPs in these four propagations (**Fig. 6D**). It appears that these six cCAPs are visited significantly more during most cCAP 8→7 and 7→8 than during contrasting propagations (*p<0.01, corrected*). Moreover, cCAPs 1, 3, and 4 show similar occurrences between two contrasting propagations, while cCAPs 2, 5, 6 show different patterns. High occurrences of cCAP 2/6 during cCAP 8→8 and cCAP 5 during cCAP 7→7 once again confirm the affinities of cCAP 2/6 toward cCAP 8 and cCAP 5 toward cCAP 7 as in **Figs. 2-4**. On the other hand, significantly lowered occurrences of cCAP 5 during cCAP 8→8 and cCAP 2/6 during cCAP 7→7 (*p< 0.001, corrected*) from the average level during contrasting propagations suggest that these lowered occurrences are the reasons behind no propagation through cCAPs 7 and 8. When these occurrence rate data are broken down according to the number of different states visited (**Fig. 6E**), significantly lowered occurrences of cCAP 5 in cCAP 8→8 and cCAP 2/6 in cCAP 7→7 (at least *p<0.05, corrected*) than other propagations in all conditions are similarly observed. No other cCAPs show consistently significant different occurrences over five different numbers of different states visited (**Supplementary Fig. 6**).

Propagation data based on 27 pairs of *pseudo-*polarized states (**Supplementary Note 4**) reveal no similar occurrence data (**Supplementary Fig. 7**), duration data (**Supplementary Figs. 8-9**), state visit data (**Supplementary Figs. 10-11**), and state-specific occurrence data (**Supplementary Fig. 12**). The propagations between the two global co-(de)activation states were unique.

### Reproducibility

We repeated the exact same analyses on other two datasets, which were independently collected (i.e., Datasets 1 and 2), and reproduced almost all phenomena reported from Dataset 0 (see dataset information in **Methods** and **Supplementary Note 1**). These phenomena include a set of cCAPs, a pair of polarized cCAPs with global co-(de)activation patterns, anti-state cCAP pairs (**Fig. 7A**), recurring patterns and their differences among different cCAPs (especially the significantly lower occurrences in two polarized cCAPs, **Fig. 7C**), oscillations at <1Hz, 5Hz, and 10Hz (data not shown), one-step transition patterns (data not shown), and propagation patterns coordinated by two polarized cCAPs (**Fig. 7B-D**). The only exception is the <1Hz oscillation, which was detected in both Datasets 0 and 1, but not obvious in Dataset 2 (having the oldest participants out of three datasets). It is important to note that the superstructure among the set of cCAPs is identified in both Datasets 1 and 2. In both datasets, this superstructure is spatially supported by the distance map (**Fig. 7A**), where brain states closer to two polarized cCAPs are also identified similarly as in Dataset 0 (i.e., cCAPs 1 and 5 close to the polarized cCAP 2, and cCAP 6 close to the polarized cCAP 7 in Dataset 1; cCAP 1 close to the polarized cCAP 2 and cCAP 4 close to the polarized cCAP 8 in Dataset 2), revealing the fine spatial constructs within eight cCAPs beyond two polarized cCAPs. The superstructure and its fine spatial constructs are then supported by one-step transition data (not shown) as in Dataset 0 (see **Fig. 4**) and propagation patterns coordinated by two polarized cCAPs (**Fig. 7D**).

**Figure 7.**
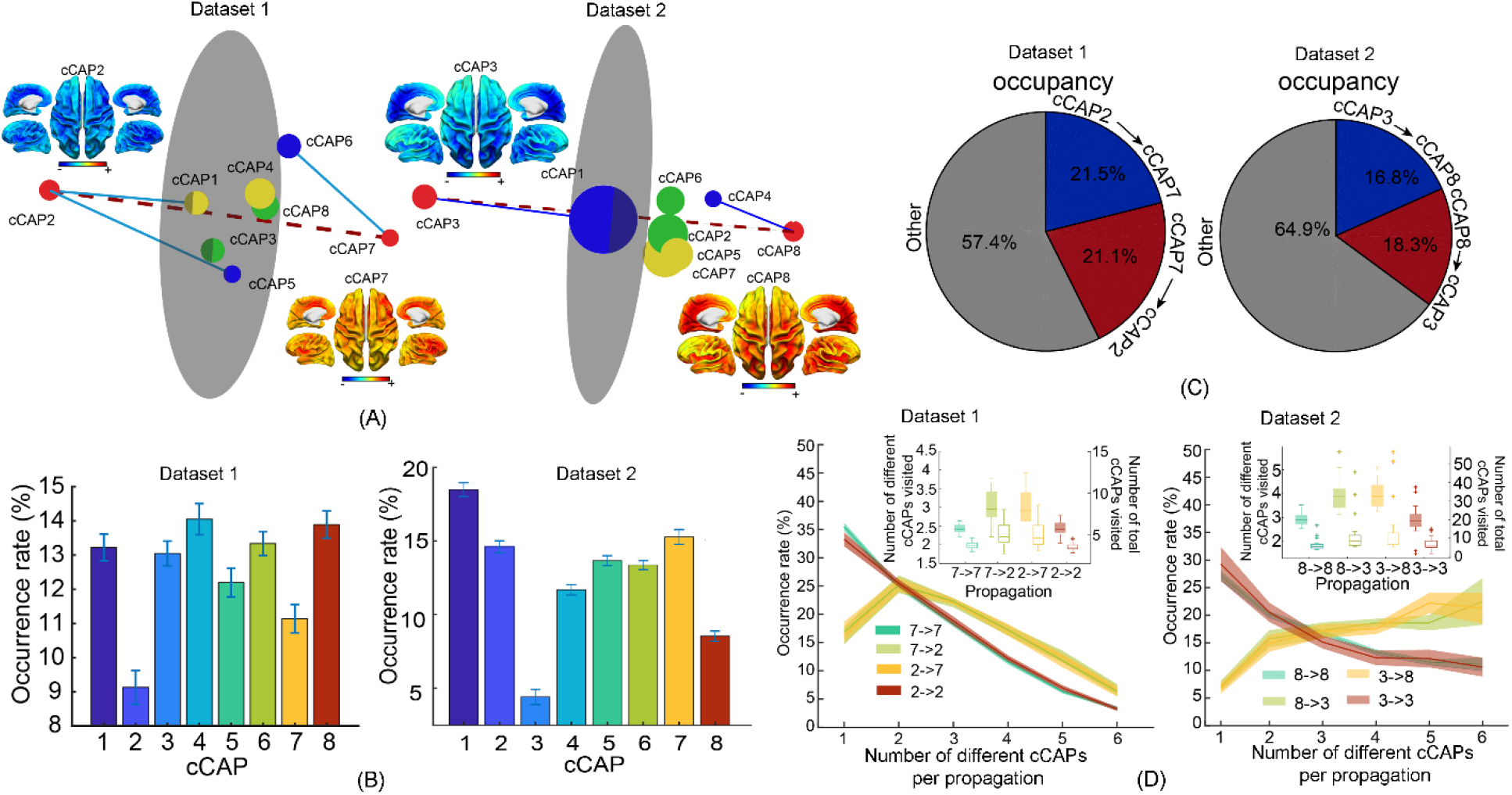
Reproduced key results of brain-wide functional states and their spatial, temporal, dynamic patterns from Datasets 1 and 2. (A) Spatial maps of two polarized global co-(de)activation brain states, i.e., cCAPs 2 and 7 in Dataset 1 and cCAPs 3 and 8 in Dataset 2, and the 3D distance maps among all cCAPs. Same-color dots: anti-state pairs; red dots: two polarized states connected by the dashed line; blue lines: connecting brain states that are structurally closest to two polarized states; gray circular plane: halfway between two polarized states. (see Fig. 2). (B) Occurrence rates of cCAPs (see Fig. 3). (C) Occupancy data of the propagations between two polarized brain states (see Fig. 4). (D) Occurrence rates of four types of propagations as a function of the number of cCAPs visited per propagation (see Fig. 6 for more details).

## Discussion

Using an advanced computational framework, we reconstructed and visualized spatial maps of brain-wide intrinsic neural networks and their dynamics in resting human brains from electrophysiological signals. Our results collectively indicate that spontaneous human brain activity is a nonstationary phenomenon, involving reconfiguration into recurring and propagating functional states, which replicate and extend previous discoveries from optical imaging studies in animals^28,29^. Such recurring and propagating spontaneous neural activity results in synchronous neural co-(de)activations across hemispherically-symmetric and functionally-connected brain areas, including the prominent phenomena of global patterns^30,35^ and anti-state pairs of co-(de)activations^25,27,28^ in their spatial topographies. This view is further expanded by reporting that time-varying patterns of spontaneous resting-state EEG signals are governed by a limited number of functional states showing rich organized dynamic structures across a wide frequency band, i.e., multi-scale oscillations from fast (5Hz and 10Hz) to slow (0.6Hz) rates. In parallel to oscillatory structures, we further found a superstructure among the identified set of functional states that regulates between-state propagations mediated by two unique states with opposite global co-(de)activation spatial patterns.

These findings advance our understanding of the principles underlying spontaneous neural networks in multiple directions. First, our results directly visualize large-scale brain-wide intrinsic functional networks based on electrical signals generated by the human brain, which have only been reported in hemodynamic signals in human^24,27,35^. Such visual constructs confirm hypothesized brain-wide networked dynamics based on electrical recordings from a limited number brain sites^20^. As several spatial prominent features of identified functional states resemble those from recent reports of human fMRI data^27,35^, our greater temporal resolution of electrophysiological recordings extends those observations and provides novel mechanistic understanding of neural determinants of brain-wide hemodynamic structures. Second, the use of signal amplitude as the basis for clustering similarities and identifying differences among moment-to-moment cortical activation topographies provides a means to discover brain-wide co-activation patterns in population-level electrical signals. In contrast, correlation-based methods applied to resting-state EEG and magnetoencephalography (MEG) signals have led to largely regionally activated neural patterns^2,33,34^ compared to distributed network patterns seen in corresponding correlation-based fMRI studies. Finally, in contrast to the slow response in hemodynamic signals^12,13^ to be <0.1Hz^28^, our present results have revealed rich frequency-specific phenomena in the classical range of EEG between 0.1Hz and 50Hz. Future studies can expand observations on both ends of the spectrum (i.e., <0.1Hz and >50Hz) to investigate more frequency-specific neuronal communications as well as cross-frequency mechanisms^36^.

Our results indicate that the number of functional states (i.e., 8) that explain most resting-state EEG temporal dynamics is considerably lower than the common number of RSNs (e.g., 15-50) identified as spatial independent sources (ICs) using ICA from resting-state fMRI^37^. Meanwhile, the dynamic states arising from these spatial sources have been suggested to be typically small (e.g., 4-8) in resting-state fMRI from both humans^38^ and animals^28^. Our eight recurring functional states (**Fig. 2**) are in fact constructed from 14 RSN components (**Supplementary Fig. 1**). This is consistent with the suggestion that ICs present a spatial parcellation of the brain rather than distinct states of functional connectivity^39^, which has been used to identify function-based parcellations of the human brain^40^. Therefore, our observations suggest, together with previous studies on dynamic states^33^, that, under resting conditions, most of these spatial independent sources may be concomitantly engaged in coordinated patterns of co-(de)activations. Co-(de)activations of these distinct spatial sources then inevitably generate brain-wide cortical patterns indicating involvement of distributed neural network systems that generate moment-to-moment dynamics as suggested by fMRI data^4,24^, which reveal phenomena that have or have not been revealed via exploring data correlation structures as discussed below.

First, the identified states exhibit brain-wide co-activations both across anatomically homologous areas between hemispheres and across functional brain regions that are spatially separated, e.g., along the anterior-posterior direction. Strong hemisphere-mirrored symmetries (observed in cCAPs 1, 3, 4, 5, 7, 8, and the cCAP 2-6 pair) are the hallmark of RSNs^8^ and resting-state functional connectivity in fMRI^28^. Electrophysiological and optical studies have also suggested such hemispheric symmetries during the propagation of brain waves in both animals^15^ and humans^41^. The pattern in cCAP 2 recapitulates the cardinal feature of DMN organization from the anterior brain, i.e., medial prefrontal cortex, to the posterior brain, i.e., posterior cingulate cortex and temporo-parietal junction^42^. These anterior-posterior structured activations, together with temporal lobe activations, reflect a full distributed membership of cortical nodes for DMN as established in human fMRI data. Similar DMN patterns have been revealed in human MEG data but with missing posterior nodes using spatial ICA^34^ or more lateralized distributions using seed-based correlation methods^2,41^. As spatial ICs are more relevant to anatomic parcellations and correlation methods are stationary, the differences in characterizing DMN of our present results and previous EEG/MEG studies support the prevailing notion of time-varying functional connectivity across brain regions reported in previous studies^24,26,28,38^. Second, the opposing co-activations between DMN and various constituents of TPN revealed in cCAPs 1, 2, 3, 4, and 5 (**Fig. 2B**) corroborate the presence of a tight inverse coupling between these two systems^43^. The widely reported brain-wide anticorrelated functional networks characterized using hemodynamic signals^4^ are believed to be manifestations of this inverse coupling at the spatial scale of the entire cortex, while no studies, to the best of our knowledge, have reported their brain-wide presence in neural electrical signals. This phenomenon supports the notion that large-scale hemodynamic correlation-based network structures are converted from large-scale spatiotemporal dynamics of spontaneous neural activity^44^. Finally, global co-activations as indicated in cCAPs 7 and 8 in our results have only been reported when human hemodynamic signals are analyzed beyond the correlation structure^30,35^. For example, the presence of strong correlated DMN and TPN, that usually lead to global co-activation patterns, has been discovered in human hemodynamic signals under the influence of global signal^45^ using a quasiperiodic pattern searching algorithm^35^. Transient global co-activations have been observed in searching globally propagating waves in both hemodynamic and neural signals from rodents^29^. Furthermore, slow oscillations (<1Hz) of membrane potential are characterized with global traveling waves of both depolarizing and hyperpolarizing components^15^, which support the existence of transient global co-activations with opposite patterns.

Lifetimes and interval times between visits of these short-lived brain states exhibit large value ranges (**Fig. 3B-C**), which suggest that cortical networks in wakefulness are predominantly asynchronous. As a matter of fact, awake behaving states are traditionally called “desynchronized EEG” in contrast to large-amplitude slow oscillations observed in quiet sleep^46^. At the same time, weak and selective coherences are present with evidence of both slow (0.6Hz) and fast oscillations (5Hz and 10Hz). Frequency-specific oscillatory synchronizations are essential mechanisms for efficient neuronal coordination across the entire brain^47^. Both 5Hz and 10Hz oscillations have been well established in EEG literature as theta^48^ and alpha rhythms^49^. The <1Hz oscillation has been discovered as an overriding EEG pattern during non-REM sleep^16,19^, while recent experimental evidence indicates that slow-wave-like activity is also present in awake animals^50^. These intermittently recurring oscillations are similarly manifested in all identified brain states although their occurrence patterns (e.g., occurrence rates) indicate statistically significant differences. Therefore, our observations suggest, on top of largely nonstationary dynamics, the presence of rich multi-scale ongoing intrinsic neural rhythms under awake resting conditions, which are independent from each other and not constrained toward specific brain states.

One of the most intriguing and novel findings of our present study is the identification of a superstructure that governs the spatial, temporal, and propagating relationships among the identified brain states, which describes an important aspect in the dynamic control of states of brain function under wakeful resting conditions. The superstructure, built with two polarized states and six intermediate states, is first established via visualizing spatial distances (**Fig. 1C**) among these states and then supported by the occurrence data, which indicate statistically significantly fewer visits to two polarized states (**Fig. 2A**). Single-step transition data (**Fig. 5**) further suggest close-to-zero direct transitions between two polarized states. Finally, propagations between two polarized states, mediated by other states, take statistically longer times (**Fig. 6A**) and more visits to intermediate states (**Fig. 6C**) than no propagations. These consistent data from multiple different facets support the genuine existence of such a superstructure, which have been replicated in two independently collected datasets (**Fig. 7**), and no structured propagation patterns exist in simulations using pseudo-polarized states (**Supplementary** **Figs. 7-12**). The generation of the superstructure is believed to be driven by structured brain-wide propagations among identified brain states coordinated by two polarized states with global co-(de)activation patterns. Beyond them, six intermediate states further exhibit layered structures in which some states are more affined to two polarized states (i.e., cCAP 5 to cCAP 8; cCAP 2/6 for cCAP 7) than others in terms of both spatial distance and transition probability.

The cellular mechanism and physiological significance of this superstructure of co-activations remains an open question. Our results indicate many similarities between the identified superstructure and the slow-wave oscillation of membrane potentials^16^, including their propagational nature, occurring frequency, and correspondence between the pairs of global co-activations and the depolarizing and hyperpolarizing components of slow-wave oscillations. However, there are still significant gaps of knowledge in how these cellular neuronal phenomena are manifested in the phase-locked behaviors of neural populations recorded in EEG^51^ and how neuronal propagating waves recorded at spatially discrete locations^52^ are converted into structured dynamics of brain-wide spatial patterns. Actually, it is also possible that other types of neuronal processes underlie such structured co-activations, such as, massive activations of cortical regions during episodic high-frequency field-potential oscillations in hippocampus^53^ and potential neural activity associated with certain conscious processes, e.g., mind wandering, occurred during wakefulness. As animal studies^29,54^ have convincingly linked the propagations among brain-wide functional states from concurrent hemodynamic and neural recordings, future concurrent studies of hemodynamic and neural signals in humans to directly study such a linkage^22^ might provide useful evidence on the cellular mechanisms of the identified superstructure. Regarding its physiological significance, a close relationship to RSNs and slow-wave oscillations may already indicate its importance to understanding brain-wide memory consolidation^55^. As several fMRI studies have reported functional connectivity changes after task learning^56^ and the relationship between global co-activations and global BOLD signals, it is of great interest for future works to study how behavioral context influences the dynamics of these structured co-activations.

A methodological advancement that needs further innovative ideas is how to best compare tomographic, dynamic, and propagational co-activation patterns across different datasets, different conditions, and/or different brain signals. Leveraging the reproducibility of our identified co-activations, we have demonstrated its detections in three independent EEG datasets. However, linking co-activation patterns identified from different brain signals, e.g., EEG and fMRI, will require other algorithms. For example, the clustering algorithm used in our present study assumes all snapshots from individual time points belong to one of eight clusters, which is different from assumptions made in searching algorithms for transient events in fMRI^27,39^. The adoption of our current algorithm is due to the noisy nature of EEG recordings, the complexity of computational processes in reconstructing brain-wide co-activations (see **Methods**), and the potential of much more complicated propagation patterns in humans compared to small animals^57^. The important aspects of future research are to develop more advanced computational processes on potentially less noisy data from new sensors^58,59^ to perform such comparisons and design analytical approaches accordingly.

## Data availability

The data that support the findings of this study are available on request from the corresponding author (L.D.) through a data use agreement. The data are not publicly available due to them containing information that could compromise research participant privacy or consent. EEG preprocessing was performed using EEGLAB toolbox (https://eeglab.org) and FASTER plugin (https://sourceforge.net/projects/faster/). The segmentation and modeling were performed using FREESURFER (https://surfer.nmr.mgh.harvard.edu). Clustering analysis was conducted using the MATLAB kmeans function (https://www.mathworks.com/help/stats/kmeans.html). Group independent component analysis was performed using Fourier ICA code (https://www.cs.helsinki.fi/group/neuroinf/code/fourierica/html/fourierica.html). Codes for minimum-norm estimate in cortical source imaging and regression analysis in statistical regression tomography were implemented using MATLAB and are available from the corresponding author on reasonable request.

## Acknowledgement

We gratefully acknowledge the financial support from NSF RII Track-2 FEC 1539068 and NIH U54HD104461. The authors are grateful to Matthew W Mosconi, Jun Wang, Lauren Ethridge, Diamond Gleghorn, Benjamin C. Doudican for assistance in data collection.

## Reference

1. Leopold, D.A., Y. Murayama, and N.K. Logothetis, Very slow activity fluctuations in monkey visual cortex: implications for functional brain imaging. Cereb Cortex, 2003. 13(4): p. 422–33.

2. Hipp, J.F., et al., Large-scale cortical correlation structure of spontaneous oscillatory activity. Nat Neurosci, 2012. 15(6): p. 884–90.

3. Biswal, B., et al., Functional connectivity in the motor cortex of resting human brain using echo-planar MRI. Magn Reson Med, 1995. 34: p. 537–541.

4. Fox, M.D., et al., The human brain is intrinsically organized into dynamic, anticorrelated functional networks. Proc Natl Acad Sci U S A, 2005. 102(27): p. 9673–8.

5. Arieli, A., et al., Dynamics of ongoing activity: explanation of the large variability in evoked cortical responses. Science, 1996. 273(5283): p. 1868–71.

6. Fox, M.D., et al., Spontaneous neuronal activity distinguishes human dorsal and ventral attention systems. Proc Natl Acad Sci U S A, 2006. 103(26): p. 10046–51.

7. Larson-Prior, L.J., et al., Cortical network functional connectivity in the descent to sleep. Proc Natl Acad Sci U S A, 2009. 106(11): p. 4489–94.

8. Smith, S.M., et al., Correspondence of the brain’s functional architecture during activation and rest. Proc Natl Acad Sci U S A, 2009. 106(31): p. 13040–5.

9. Greicius, M., Resting-state functional connectivity in neuropsychiatric disorders. Curr Opin Neurol, 2008. 21(4): p. 424–30.

10. Fox, M.D. and M.E. Raichle, Spontaneous fluctuations in brain activity observed with functional magnetic resonance imaging. Nat Rev Neurosci, 2007. 8(9): p. 700–11.

11. Logothetis, N.K., What we can do and what we cannot do with fMRI. Nature, 2008. 453: p. 869–878.

12. Logothetis, N.K., et al., Neurophysiological investigation of the basis of the fMRI signal. Nature, 2001. 412(6843): p. 150–7.

13. Shmuel, A. and D.A. Leopold, Neuronal correlates of spontaneous fluctuations in fMRI signals in monkey visual cortex: Implications for functional connectivity at rest. Hum Brain Mapp, 2008. 29(7): p. 751–61.

14. Huang, X., et al., Spiral wave dynamics in neocortex. Neuron, 2010. 68(5): p. 978–990.

15. Stroh, A., et al., Making waves: initiation and propagation of corticothalamic Ca2+ waves in vivo. Neuron, 2013. 77(6): p. 1136–50.

16. Steriade, M., et al., The slow (< 1 Hz) oscillation in reticular thalamic and thalamocortical neurons: scenario of sleep rhythm generation in interacting thalamic and neocortical networks. J Neurosci, 1993. 13(8): p. 3284–99.

17. Luczak, A., et al., Sequential structure of neocortical spontaneous activity in vivo. Proc Natl Acad Sci U S A, 2007. 104(1): p. 347–52.

18. Ferezou, I., et al., Spatiotemporal dynamics of cortical sensorimotor integration in behaving mice. Neuron, 2007. 56(5): p. 907–23.

19. Crunelli, V. and S.W. Hughes, The slow (<1Hz) rhythm of non-REM sleep: a dialogue between three cardinal oscillators. nat Neurosci, 2010. 13: p. 9–17.

20. Massimini, M., et al., The Sleep Slow Oscillation as a Traveling Wave. J Neurosci, 2004. 24(31): p. 6862–6870.

21. Ritter, P. and A. Villringer, Simultaneous EEG-fMRI. Neurosci Biobehav Rev, 2006. 30(6): p. 823–38.

22. Yuan, H., et al., Reconstructing Large-Scale Brain Resting-State Networks from High-Resolution EEG: Spatial and Temporal Comparisons with fMRI. Brain Connect, 2016. 6(2): p. 122–35.

23. Feige, B., et al., Cortical and subcortical correlates of electroencephalographic alpha rhythm modulation. J Neurophysiol, 2005. 93(5): p. 2864–72.

24. Allen, E.A., et al., Tracking Whole-Brain Connectivity Dynamics in the Resting State. Cerebral Cortex, 2014. 24(3): p. 663–676.

25. Liu, X. and J.H. Duyn, Time-varying functional network information extracted from brief instances of spontaneous brain activity. Proc Natl Acad Sci U S A, 2013. 110(11): p. 4392–7.

26. Liu, X., et al., Subcortical evidence for a contribution of arousal to fMRI studies of brain activity. Nat Commun, 2018. 9(1): p. 395.

27. Karahanoglu, F.I. and D. Van De Ville, Transient brain activity disentangles fMRI resting-state dynamics in terms of spatially and temporally overlapping networks. Nat Commun, 2015. 6: p. 7751.

28. Gutierrez-Barragan, D., et al., Infraslow State Fluctuations Govern Spontaneous fMRI Network Dynamics. Curr Biol, 2019. 29(14): p. 2295–2306 e5.

29. Matsui, T., T. Murakami, and K. Ohki, Transient neuronal coactivations embedded in globally propagating waves underlie resting-state functional connectivity. Proc Natl Acad Sci U S A, 2016. 113(23): p. 6556–61.

30. Mitra, A., et al., Lag threads organize the brain’s intrinsic activity. Proc Natl Acad Sci U S A, 2015. 112(17): p. E2235–E2244.

31. Grienberger, C. and A. Konnerth, Imaging calcium in neurons. Neuron, 2012. 73(5): p. 862–85.

32. Fischl, B., FreeSurfer. Neuroimage, 2012. 62(2): p. 774–81.

33. Shou, G., et al., Whole-brain electrophysiological functional connectivity dynamics in resting-state EEG. J Neural Eng, 2020. 17(2): p. 026016.

34. Brookes, M.J., et al., Investigating the electrophysiological basis of resting state networks using magnetoencephalography. Proc Natl Acad Sci U S A, 2011. 108(40): p. 16783–8.

35. Yousefi, B., et al., Quasi-periodic patterns of intrinsic brain activity in individuals and their relationship to global signal. Neuroimage, 2018. 167: p. 297–308.

36. Canolty, R.T. and R.T. Knight, The functional role of cross-frequency coupling. Trends Cogn Sci, 2010. 14(11): p. 506–15.

37. Damoiseaux, J.S., et al., Consistent resting-state networks across healthy subjects. Proc Natl Acad Sci U S A, 2006. 103(37): p. 13848–53.

38. Calhoun, V.D., et al., The chronnectome: time-varying connectivity networks as the next frontier in fMRI data discovery. Neuron, 2014. 84(2): p. 262–274.

39. Liu, X., C. Chang, and J.H. Duyn, Decomposition of spontaneous brain activity into distinct fMRI co-activation patterns. Front Syst Neurosci, 2013. 7: p. 101.

40. Smith, S.M., et al., Functional connectomics from resting-state fMRI. Trends Cogn Sci, 2013. 17(12): p. 666–82.

41. de Pasquale, F., et al., Temporal dynamics of spontaneous MEG activity in brain networks. Proc Natl Acad Sci U S A, 2010. 107(13): p. 6040–5.

42. Buckner, R.L., J.R. Andrews-Hanna, and D.L. Schacter, The brain’s default network: anatomy, function, and relevance to disease. Ann N Y Acad Sci, 2008. 1124: p. 1–38.

43. Popa, D., A.T. Popescu, and D. Pare, Contrasting activity profile of two distributed cortical networks as a function of attentional demands. J neurosci, 2009. 29(4): p. 1191–1201.

44. Tagliazucchi, E., et al., Criticality in large-scale brain FMRI dynamics unveiled by a novel point process analysis. Front Physiol, 2012. 3: p. 15.

45. Liu, T.T., A. Nalci, and M. Falahpour, The global signal in fMRI: Nuisance or Information? Neuroimage, 2017. 150: p. 213–229.

46. Steriade, M., D.A. McCormick, and T.J. Sejnowski, Thalamocortical oscillations in the sleeping and aroused brain. Science, 1993. 262(5134): p. 679–85.

47. Siegel, M., T.H. Donner, and A.K. Engel, Spectral fingerprints of large-scale neuronal interactions. Nat Rev Neurosci, 2012. 13(2): p. 121–34.

48. Kahana, M.J., et al., Human theta oscillations exhibit task dependence during virtual maze navigation. Nature, 1999. 399(6738): p. 781–4.

49. Halgren, M., et al., The generation and propagation of the human alpha rhythm. Proc Natl Acad Sci U S A, 2019. 116(47): p. 23772–23782.

50. Vyazovskiy, V.V., et al., Local sleep in awake rats. Nature, 2011. 472(7344): p. 443–7.

51. Wang, X.J., Neurophysiological and computational principles of cortical rhythms in cognition. Physiol Rev, 2010. 90(3): p. 1195–268.

52. He, B.J., et al., Electrophysiological correlates of the brain’s intrinsic large-scale functional architecture. Proc Natl Acad Sci U S A, 2008. 105(41): p. 16039–44.

53. Logothetis, N.K., et al., Hippocampal-cortical interaction during periods of subcortical silence. Nature, 2012. 491 (7425): p. 547–53.

54. Schwalm, M., et al., Cortex-wide BOLD fMRI activity reflects locally-recorded slow oscillation-associated calcium waves. Elife, 2017. 6.

55. Tambini, A., N. Ketz, and L. Davachi, Enhanced brain correlations during rest are related to memory for recent experiences. Neuron, 2010. 65(2): p. 280–90.

56. Lewis, C.M., et al., Learning sculpts the spontaneous activity of the resting human brain. Proc Natl Acad Sci U S A, 2009. 106(41): p. 17558–63.

57. Mitra, A. and M.E. Raichle, How networks communicate: propagation patterns in spontaneous brain activity. Philos Trans R Soc Lond B Biol Sci., 2016. 371(1705): p. 20150546.

58. Aghaei-Lasboo, A., et al., Tripolar concentric EEG electrodes reduce noise. Clin Neurophysiol, 2020. 131(1): p. 193–198.

59. Boto, E., et al., Moving magnetoencephalography towards real-world applications with a wearable system. Nature, 2018. 555(7698): p. 657–661.

60. Shou, G., et al., Electrophysiological signatures of atypical intrinsic brain connectivity networks in autism. J Neural Eng, 2017. 14(4): p. 046010.

61. Richards, J.E., et al., A database of age-appropriate average MRI templates. Neuroimage, 2016. 124(Pt B): p. 1254–1259.

62. Ding, L., et al., Lasting modulation effects of rTMS on neural activity and connectivity as revealed by resting-state EEG. IEEE Trans Biomed Eng, 2014. 61(7): p. 2070–80.

63. Delorme, A. and S. Makeig, EEGLAB: an open source toolbox for analysis of single-trial EEG dynamics including independent component analysis. J Neurosci Methods, 2004. 134(1): p. 9–21.

64. Lai, Y., et al., Estimation of in vivo human brain-to-skull conductivity ratio from simultaneous extra- and intra-cranial electrical potential recordings. Clinical Neurophysiology, 2005. 116(2): p. 456–465.

65. Hamalainen, M.S. and J. Sarvas, Realistic Conductivity Geometry Model of the Human Head for Interpretation of Neuromagnetic Data. Ieee Transactions on Biomedical Engineering, 1989. 36(2): p. 165–171.

66. Hamalainen, M.S. and R.J. Ilmoniemi, Interpreting Magnetic-Fields of the Brain - Minimum Norm Estimates. Medical & Biological Engineering & Computing, 1994. 32(1): p. 35–42.

67. Golub, G.H., M. Heath, and G. Wahba, Generalized Cross-Validation as a Method for Choosing a Good Ridge Parameter. Technometrics, 1979. 21(2): p. 215–223.

68. Bingham, E. and A. Hyvarinen, A fast fixed-point algorithm for independent component analysis of complex valued signals. Int J Neural Syst, 2000. 10(1): p. 1–8.

69. Shou, G.F., L. Ding, and D. Dasari, Probing neural activations from continuous EEG in a real-world task: Time-frequency independent component analysis. Journal of Neuroscience Methods, 2012. 209(1): p. 22–34.

70. Woolrich, M.W., et al., Temporal autocorrelation in univariate linear modeling of FMRI data. Neuroimage, 2001. 14(6): p. 1370–86.

