## supplemental file for "Brain-wide neural co-activations in resting human"

Ding et al.

### Supplementary Note 1: Dataset 1 and Dataset 2

Dataset 1: The dataset was obtained from healthy controls in our previous experiments that were recruited for examining EEG signatures to be compared with patients of autism spectrum disorders (ASD)[1]. The study was approved by the local Institutional Review Board, and written informed consent was obtained according to the Declaration of Helsinki for each adult participant and informed parental consent was obtained for individuals less than 18 years old. Resting-state EEG data with eye-closed were recorded for 5 minutes at a sample frequency of 512Hz using a Biosemi ActiveTwo 128-channel 24-bit resolution system, with active electrodes and low-passed with a 5$^{th}$ order digital sync anti-aliasing filter. During the recording, participants sat on a reclining chair in a sound attenuated room with a dim light and were monitored for vigilance. For this dataset, there was no structure MRI recording from individuals. Instead, age-appropriate MRI templates[2] were used.

Dataset 2: This dataset is obtained from healthy controls in our previous experiment that were recruited for examining EEG signatures to be compared with patients of a balance disorder[3,4]. The study was approved by Western IRB, and written informed consent was obtained from all participants before the start of all procedures. Five minutes resting-state EEG data with eye-closed were recorded at a sample frequency of 1000Hz using a 126-channel BrainAmp amplifier (Brain Products GmbH, Munich, Germany). During the recording, participants were sitting in a recliner quietly in a darkened room. Prior to the EEG recording, individual structural MRI data were acquired on a General Electric (GE) Discovery MR750 3 T MRI whole-body scanner (GE Healthcare, Milwaukee WI, USA), which had the parameters: FOV=240mm, axial slices per slab = 190, slice thickness = 0.9mm, image matrix = 256×256, TR/TE = 5/2.012ms, acceleration factor R=2, flip angle = 8°, inversion time TI = 725ms, and sampling bandwidth = 31.2kHz. In addition, EEG sensor positions and three landmark fiducial locations (i.e., nasion, left and right pre-auricular points) were digitized by the Polhemus Patriot system.

### Supplementary Note 2: Spatial and Temporal Patterns of cCAPs in Figs. 2-3

Regarding the results in **Fig. 2A-B**, the co-(de)activation patterns of all cCAPs could be observed at two levels: cortical maps (**Fig. 2A**) and activity levels of EEG component signals (i.e., RSNs), which are indicated in the weights of the cluster center vectors (**Fig. 2B**). Cortical maps of six cCAPs (cCAPs 1, 3, 4, 5, 7, and 8) show symmetric patterns between the two hemispheres, and the rest two cCAPs (cCAPs 2 and 6) are symmetric as a pair with opposite hemispheric dominances. The co-activations and co-deactivations of these cCAPs are further revealed at the activity level of individual RSNs (EEG component signals), which are spatially non-overlapping (**Supplementary Fig. 1B**), as the building blocks of identified cCAPs. Each cCAP (from cCAP 1 to cCAP 6) consists of at least two RSNs exhibiting the top two negative magnitudes and at least two other RSNs at the top two positive magnitudes (**Fig. 2B**) of their own individual ranges. Two additional cCAPs (cCAPs 7 and 8) indicate global co-activation and co-deactivation patterns, respectively, as all RSNs reach their own top (or close to top) levels of positive or negative magnitude of activity, which are further supported by their spatial maps (**Fig. 2A**).

The superstructure formulated by the set of eight cCAPs, as illustrated in **Fig. 2C**, shows more detailed and layered spatial relationships among them beyond two polarized states (cCAPs 7 and 8). Firstly, two anti-state pairs (cCAPs 1-4 and cCAPs 3-5) are separated by the middle interface, which is half-way between cCAPs 7 and 8, as one of each pair is close to cCAP 8 and another is close to cCAP 7. Secondly, by calculating the distance ratio between each of six cCAPs toward cCAP 7 with respect to cCAP 8 (**Supplementary Fig. 2A**), cCAP 5 is closest to the polarized state of cCAP 8 while both cCAPs 2 and 6 are closest to the polarized state of cCAP 7. The fact indicates that, structurally, cCAP 5 is more affined toward cCAP 8, while cCAPs 2 and 6 are more affined toward cCAP 7, which are also supported by temporal transitional and propagational data (**Figs. 4 and 6D**) discussed below.

Regarding the frequencies of two characteristic peaks in the histograms of cCAP interval times (**Fig. 3B**), they are estimated counting both lifetimes and interval times of corresponding cCAPs. Interestingly, all cCAPs exhibit these peaks in similar time windows, with the first one between 60 to 70ms and the second one between 170 to 180ms. Considering the mean lifetime of ~30ms for individual cCAP occurrences, after inversion of the sum of lifetime and interval time, these numbers put the first peak in the frequency of ~10Hz and the second peak in the frequency of ~5Hz.

### Supplementary Note 3: statistical analysis in Fig. 6 and supplementary Fig. 6

A series of statistical analyses were performed on different metrics (**Fig. 6**) related to the propagations between two polarized states. For the metric of occurrences (**Fig. 6A**), paired *t* tests were performed in the comparisons of two actual propagations between two polarized states (i.e., cCAP 7→8 and cCAP 8→7) and two contrasting propagations (i.e., propagations between the same polarized states, cCAP 7→7 and cCAP 8→8). To control for multiple comparisons, Bonferroni adjustment was performed and the adjusted *p* values less than 0.05 were considered to be significant. The actual *p* values after adjustment were included in the figure. The mean occurrences across all individuals for cCAP 7→7 and cCAP 8→8 were 1378 and 1503, respectively, which were >85% higher than the mean occurrences for cCAP 7→8 and cCAP 8→7, i.e., 739 for both.

For the metric of propagation durations (**Fig. 6B**), paired *t* tests were performed in the comparisons of two actual propagations and two contrasting propagations on the mean values of durations (the inset) while the boxplots with median values were used in **Fig. 6B.** To control for multiple comparisons, Bonferroni adjustment was performed and the adjusted *p* values less than 0.05 was considered to be significant. The actual *p* values after adjustment were also included in the figure. The mean propagation durations for cCAP 7→8 and cCAP 8→7 were 152 and 157ms, respectively, which are about 1.6 times longer than the mean propagation durations for cCAP 7→7 and cCAP 8→8, i.e., 95 and 99ms, respectively. For the histogram data of propagation durations, statistical analyses were only performed on the duration values with the number of samples (i.e., participants) larger than 10 (**Supplementary Fig. 5**) as any of four types of propagations might have zero occurrences in some participants for specific durations. Repeated measures ANOVA (rmANOVA) were first performed among data from the four types of propagations at each duration, respectively, and *post-hoc t* tests were performed on the durations that had significant *p* values in the rmANOVA tests ($p < 0.01$ and consecutive durations > 5). Basically, the significant differences were considered when *p* values were less than 0.01 on at least five consecutive duration values.

For the metrics related to the numbers of cCAPs visited per propagation (including both numbers of total cCAPs visited and different cCAPs visited, **Fig. 6C**), paired *t* tests were performed for all possible pairs (i.e., 6 pairs) of different propagations (the inset). Bonferroni adjustment was performed and the adjusted *p* values less than 0.05 were considered to be significant. The actual *p* values after adjustment were included in the figure. It is noted that the numbers of total cCAPs visited in cCAP 7→8 and cCAP 8→7 (the mean values are 5.51 and 5.65, respectively) are about 1.5 times more than the corresponding mean values for cCAP 7→7 and cCAP 8→8 (3.74 and 3.73, respectively). The numbers of different cCAPs visited in cCAP 7→8 and cCAP 8→7 (the mean values for different cCAPs visited are 3.11 and 3.12, respectively) were about 1.25 times more than the corresponding mean values for cCAP 7→7 and cCAP 8→8 (2.54 and 2.46, respectively). For the occurrence rates of four types of propagations as the function of the number of different cCAPs visited, rmANOVA tests were first performed at each number of different cCAPs visited (i.e., from 1 to 6). Bonferroni adjustment was performed and the adjusted *p* values less than 0.05 were claimed as significant. The actual *p* values after adjustment were included in the figure. Thereafter, *post-hoc t* tests were performed to compare the values between the actual propagations between cCAPs 7 and 8 and the contrasting propagations at the numbers of different cCAPs visited in which the significances were indicated in the rmANOVA tests. Bonferroni adjustments were performed for the total number of comparisons as the product of the number of conditions of significance in rmANOVA (i.e., 4) and the number of comparisons between two actual propagations and two contrasting propagations (i.e., 4). The adjusted *p* values less than 0.05 were claimed as significant and the adjusted *p* values were included in the figure caption.

For the metric of the occurrence rate of each cCAP within four types of propagations (**Fig. 6D**), rmANOVA tests were firstly performed on the four types of propagations for each cCAP. Bonferroni adjustment was performed and the adjusted *p* values less than 0.05 were considered significant. Thereafter, *post-hoc t* tests were performed to compare data between all possible pairs of four propagations in the cases of cCAPs of significance in the rmANOVA tests. Bonferroni adjustments were performed for the total number of comparisons as the product of the number of cCAPs of significance in rmANOVA (i.e., 6), and the number of comparisons for each cCAP (i.e., 6). The adjusted *p* values less than 0.05 were considered significant and the adjusted *p* values were included in the figure. Beyond the definitions of four types of propagational patterns coordinated by cCAPs 7 and 8 (**Fig. 6A**), the roles of other cCAPs in these propagational patterns have also distinguished them into two different groups (i.e., cCAPs 2, 5, 6 as one group and cCAPs 1, 3, 4 as another group), which further reveals detailed structures in these propagations. More specifically, for cCAPs 1, 3, and 4, their occurrences during two contrasting propagations are largely similar with no significant differences. For cCAPs 2/6, their occurrences within cCAP 7→7 are elevated close to or the same as the level in cCAP 8→7 and 7→8, while their occurrences within cCAP 8→8 are further significantly lowered (*p< 0.001, corrected*) from the average level during reference propagations. Conversely, the occurrences of cCAP 5 are elevated during cCAP 8→8 and further decreased during cCAP 7→7. These observations suggest that fewer occurrences of cCAPs 2/6 during cCAP 7→7 and fewer occurrences of cCAP 5 during cCAP 8→8 are the reasons behind no propagation through cCAPs 7 and 8. High occurrences of cCAPs 2/6 during cCAP 8→8 and cCAP 5 during cCAP 7→7 provide further evidence that cCAPs 2/6 are close to cCAP 8 and cCAP 5 is close to cCAP 7.

For the metric of the occurrence rate of each cCAP within four types of propagations as a function of the number of different cCAPs visited (**Fig. 6E**), rmANOVA tests were first performed for each cCAP on all cases of the number of different cCAPs visited, respectively. Bonferroni adjustment was performed for the total number of comparisons as the product of the number of cCAPs (i.e., 6) and the number of different cCAPs visited for each cCAP, (i.e., 5, the condition with the number of different cCAPs visited equal to 6 was not considered since the occurrence rates for all cCAPs in this condition are same). The adjusted *p* values less than 0.05 were considered significant and the adjusted *p* values were included in the figure.

Thereafter, *post-hoc t* tests were performed to compare data between all possible pairs of four propagations at each condition for the number of different cCAPs visited for each cCAP of significance in the rmANOVA tests. Bonferroni adjustments were performed for the total number of pairs of propagations (i.e., 6). The adjusted *p* values less than 0.05 were considered significant and the adjusted *p* values were included in the figure. These occurrence rate data broken down from **Fig. 6D** indicate the similar relationships that cCAPs 2/6 are close to cCAP 8 and cCAP 5 is close to cCAP 7. More specifically, cCAP 5 is consistently lower (at least $p<0.05$, *corrected*) during cCAP 7→7 than other propagations and cCAP 6 is consistently lower (at least $p<0.05$, *corrected*) and cCAP 2 is mostly lower (at least $p<0.05$, *corrected*) during cCAP 8→8 from the conditions of one to five different state(s) visited.

### Supplementary Note 4: Metrics calculated on propagations between any two *pseudo*-polarized states

To investigate whether the propagations between two polarized cCAPs (i.e., cCAPs 7 and 8) were unique, we calculated all same metrics on the propagations between two *pseudo*-polarized states simulated from all possible pairs of eight cCAPs other than the cCAPs 7-8 pair. Accordingly, outcome data for the occurrences for four types of propagations (corresponding to **Fig. 6A**), the histograms of durations for four types of propagations (corresponding to **Fig. 6B),** the mean durations for four types of propagations (corresponding to the inset of **Fig. 6B**), the occurrence rates of four types of propagations at various numbers of different cCAPs visited (corresponding to **Fig. 6C**), the numbers of both different cCAPs visited and total cCAPs visited during four types of propagation (corresponding to the inset of **Fig. 6C**), and the occurrence rates of six cCAPs other than two *pseudo*-polarized states within four types of propagations (corresponding to **Fig. 6D**), were presented in the **Supplementary Figs. 7-12**, respectively. It was noted that the propagation patterns observed between the two real polarized states (i.e., cCAPs 7 and 8) were unique and no other simulated pairs of *pseudo*-polarized states suggested similar patterns indicated by all these metrics (see more in **Supplementary Figs. 7-12**).

### Reference


1. Shou, G., et al., *Electrophysiological signatures of atypical intrinsic brain connectivity networks in autism.* J Neural Eng, 2017. **14**(4): p. 046010.
2. Richards, J.E., et al., *A database of age-appropriate average MRI templates.* Neuroimage, 2016. **124**(Pt B): p. 1254-1259.
3. Ding, L., et al., *Lasting modulation effects of rTMS on neural activity and connectivity as revealed by resting-state EEG.* IEEE Trans Biomed Eng, 2014. **61**(7): p. 2070-80.
4. Chen, Y., et al., *Multimodal Imaging of Repetitive Transcranial Magnetic Stimulation Effect on Brain Network: A Combined Electroencephalogram and Functional Magnetic Resonance Imaging Study.* Brain Connect, 2019. **9**(4): p. 311-321.

### Supplementary Figures

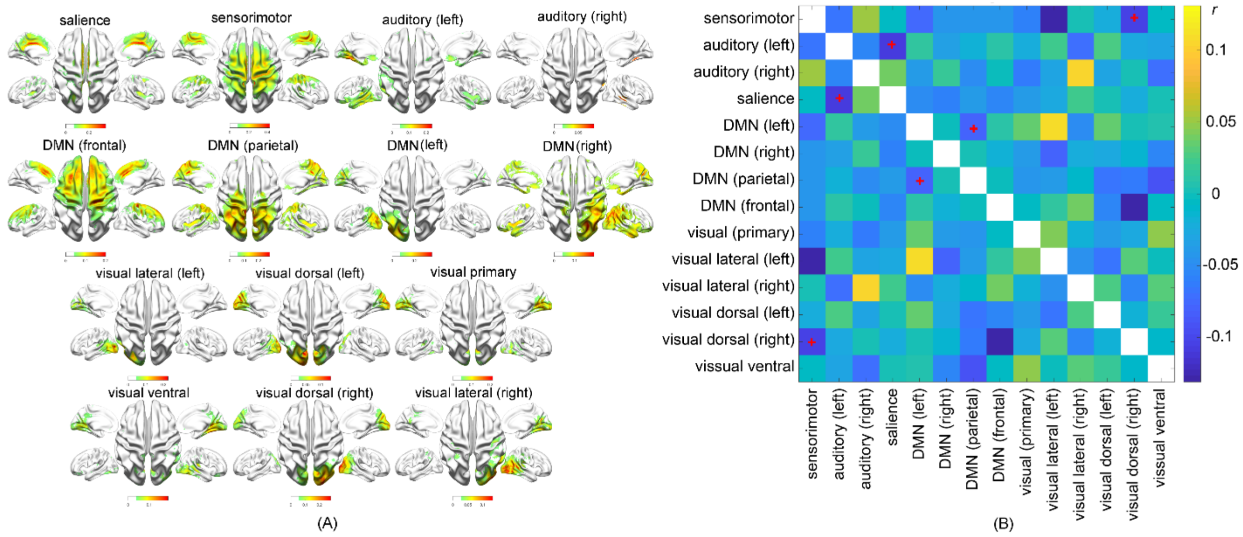


Supplementary Figure 1. (A) Cortical spatial maps of 14 selected RSNs obtained from the 1st regression in the step of statistical regression tomography (**Fig. 1A**). (B) Correlation coefficients on vectorized cortical spatial maps for any pairs of RSNs. Only 3 out of total 91 RSN pairs showed significantly anti-correlation values (denoted as '+', $p$<0.0005, FDR corrected) against the hypothesized zero spatial correlation, indicating that all identified RSNs were not confused in terms of their spatial maps.

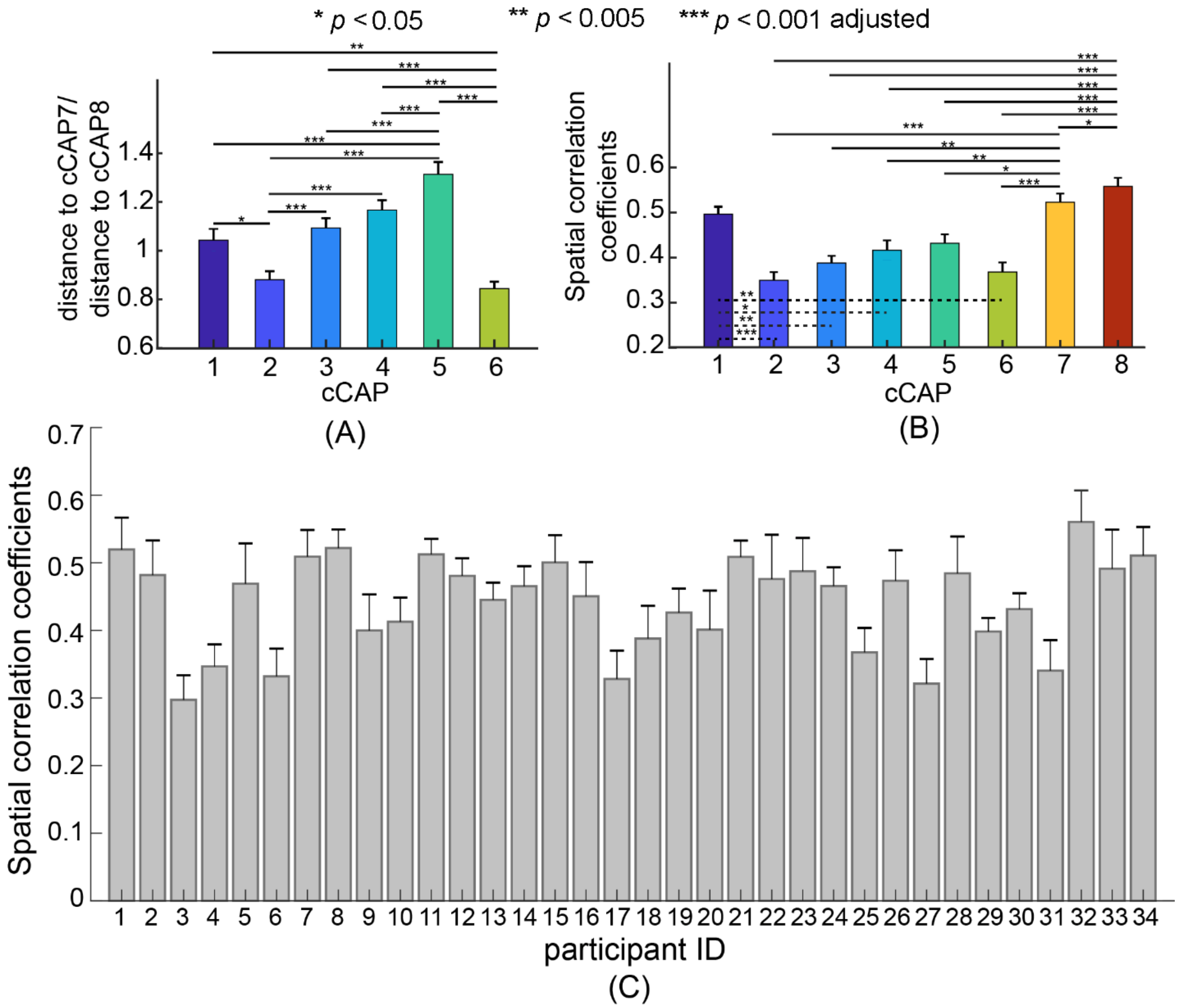


Supplementary Figure 2. (A) The ratios between the distance to cCAP 7 and the distance to cCAP 8 for the rest six cCAPs. The ratio for cCAP 5 is significantly higher than all other five cCAPs and has the value mostly different from one (equal distance to cCAP 7 and cCAP 8) toward the direction of large values, indicating that cCAP 5 is structurally closer to cCAP 7 than other five cCAPs in the set of identified functional brain states. The ratios for cCAPs 2 and 6 are significantly lower than all other four cCAPs and have the values mostly different from one toward the direction of small values, indicating that cCAPs 2 and 6 are structurally closer to cCAP 8 than other four cCAPs in the set of identified functional brain states. Here, the distances were calculated using the L1-norm metric between the cluster centers of cCAP. (B) Spatial correlation coefficients (SCC) calculated on vectorized cortical spatial maps between those from individual participants and the grand-averaged one of each cCAP, which are all statistically significantly larger than zero (at least $p$<1e-6, Bonferroni adjusted). In particular, cCAPs 7, and 8 indicate the highest reproducibility (only two SCCs larger than 0.5), and cCAP 1 also shows a high reproducibility with SCC close to 0.5. These three SCCs are statistically significantly higher than SCCs from other five cCAPs. (C) Mean ($\pm$SEM) values of spatial correlation coefficients between cCAPs from individuals and the corresponding grand-averaged ones in (B) presented for individual participants. Their mean values range between 0.3 and 0.56, which are all statistically significantly larger than zero (at least $p < 0.05$, Bonferroni adjusted). Note that the $p$ values reported in (A) and (B) were Bonferroni adjusted.

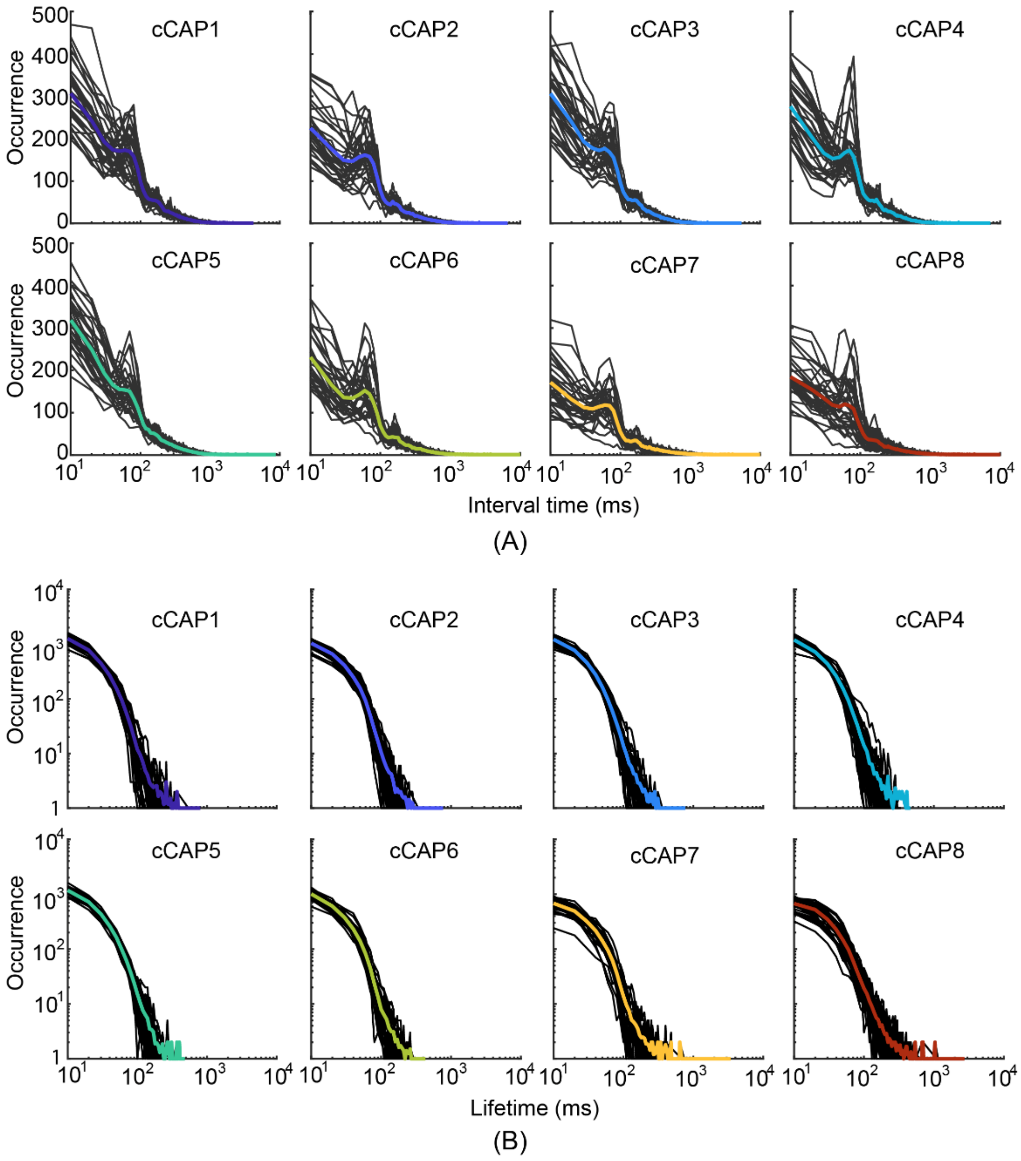


Supplementary Figure 3. (A) Histograms of interval time for all participants with each panel for one cCAP. Two peaks (the first one between 65 to 75ms and the second one between 170 to 180ms) indicating intrinsic dynamic rhythms revealed at the histograms of the group-level cCAPs could be similarly observed in the histograms of individual-level cCAPs. The fact suggests the consistencies of temporal dynamics of cCAPs between the group-level and individual-level. (B) Histograms of lifetime for all participants with each panel for one cCAP, which also indicate consistencies between the group-level and individual-level. Each thin black line indicates one participant, and the thick color one indicates the grand-averaged one across all participants.

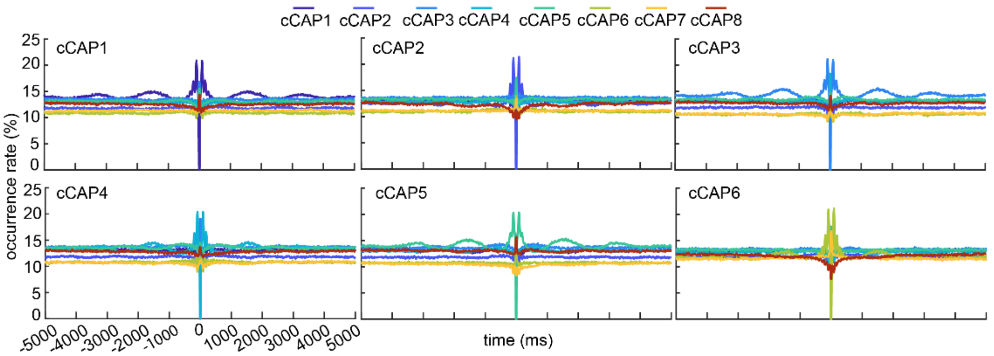
cCAP1 cCAP2 cCAP3 cCAP4 cCAP5 cCAP6 cCAP7 cCAP8
cCAP1
cCAP2
cCAP3
cCAP4
cCAP5
cCAP6
occurrence rate (%)
time (ms)
-5000 -4000 -3000 -2000 -1000 0 1000 2000 3000 4000 5000

(A)

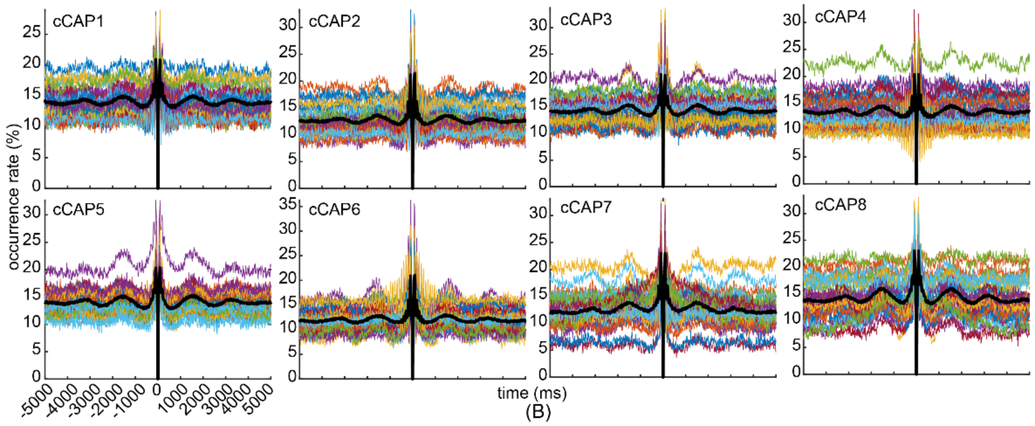
cCAP1
cCAP2
cCAP3
cCAP4
cCAP5
cCAP6
cCAP7
cCAP8
occurrence rate (%)
time (ms)
-5000 -4000 -3000 -2000 -1000 0 1000 2000 3000 4000 5000

(B)

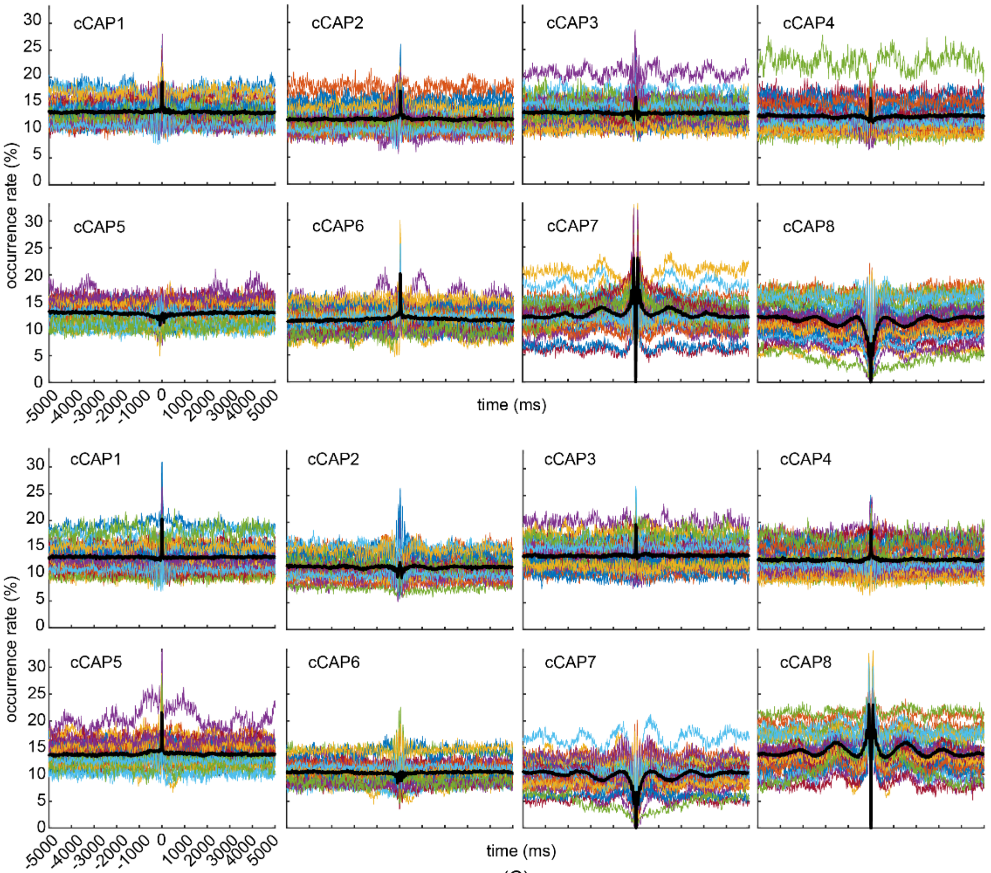
cCAP1
cCAP2
cCAP3
cCAP4
cCAP5
cCAP6
cCAP7
cCAP8
occurrence rate (%)
time (ms)
-5000 -4000 -3000 -2000 -1000 0 1000 2000 3000 4000 5000

(C)

Supplementary Figure 4. Occurrence probabilities of cCAPs as functions of time distances (between -5000 ms to 5000 ms) toward a target cCAP. (A) Group-level occurrence probabilities of all cCAPs at different time distances toward the target cCAP (marked in the top left corner of each panel, see **Fig. 3** for cCAP 7 or 8 as the target cCAP). (B) Occurrence probabilities of all cCAPs at different time distances toward the target cCAP as themselves (marked in the top left corner of each panel) both at the group-level (thick black curves) and individuals (thin color curves). (C) Occurrence probabilities of all cCAPs (marked in the top left corner of each panel) at different time distances toward the target cCAP as cCAP 7 (top) or cCAP 8 (bottom) both at the group-level (thick black curves) and individuals (thin color curves). Note that two oscillations are observed in above plots with one of short scale having the inter-peak distance of ~100 ms and another one of long scale showing the inter-peak distance of ~1.6 s. The long-scale oscillations could be observed mainly when the occurrence probabilities are calculated on cCAPs to the target cCAP as themselves (see the panel B). In addition, in the pair of cCAPs 7-8, the elevated oscillatory phenomena (for both short-scale and long-scale oscillations) in one of them lead to the symmetric but depressed oscillatory phenomena in another (see the panel C). These elevated oscillatory phenomena can be reliably detected both at the group and individual levels (see the panels B and C).

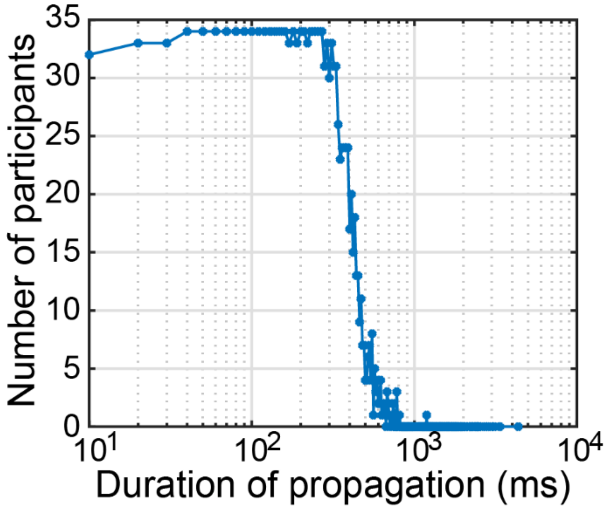


Supplementary Figure 5. The histogram of the number of participants who have the specific duration values of all four propagations: cCAP 7→7, cCAP 7→8, cCAP 8→7, and cCAP 8→8. The statistical comparisons between two propagations involving two polarized states (i.e., cCAP 7→8 and cCAP 8→7) and two contrasting propagations (i.e., cCAP 7→7 and cCAP 8→8) were only conducted for durations with the number of participants larger than 10 to maintain sufficient power for statistical analysis (see results in **Fig. 6B**).

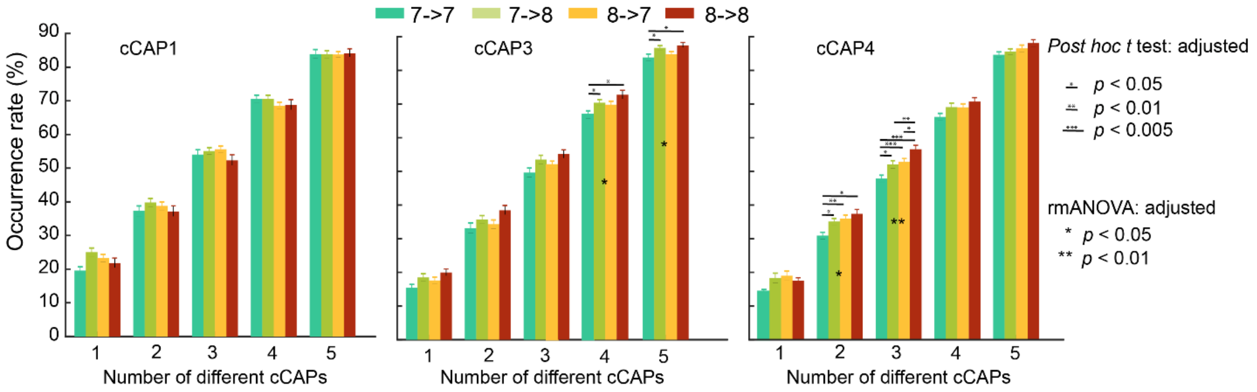


Supplementary Figure 6. Occurrence rates of cCAPs 1, 3 and 4 within each type of propagation (i.e., cCAP 7→7, cCAP 7→8, cCAP 8→8, and cCAP 8→7) as functions of number of different cCAPs visited per propagation. In comparison to cCAPs 2, 5, and 6 (see **Fig. 6E**), these cCAPs indicate much less differences in their roles in facilitating the propagations between two polarized states (i.e., cCAP 7→8 and cCAP 8→7) as compared with their roles in two contrasting propagations (i.e., cCAP 7→7 and cCAP 8→8). The condition of number of cCAPs as 6 is omitted since all occurrence rates are 100% by the definition of this metric. Note that the *p* values reported were Bonferroni adjusted.

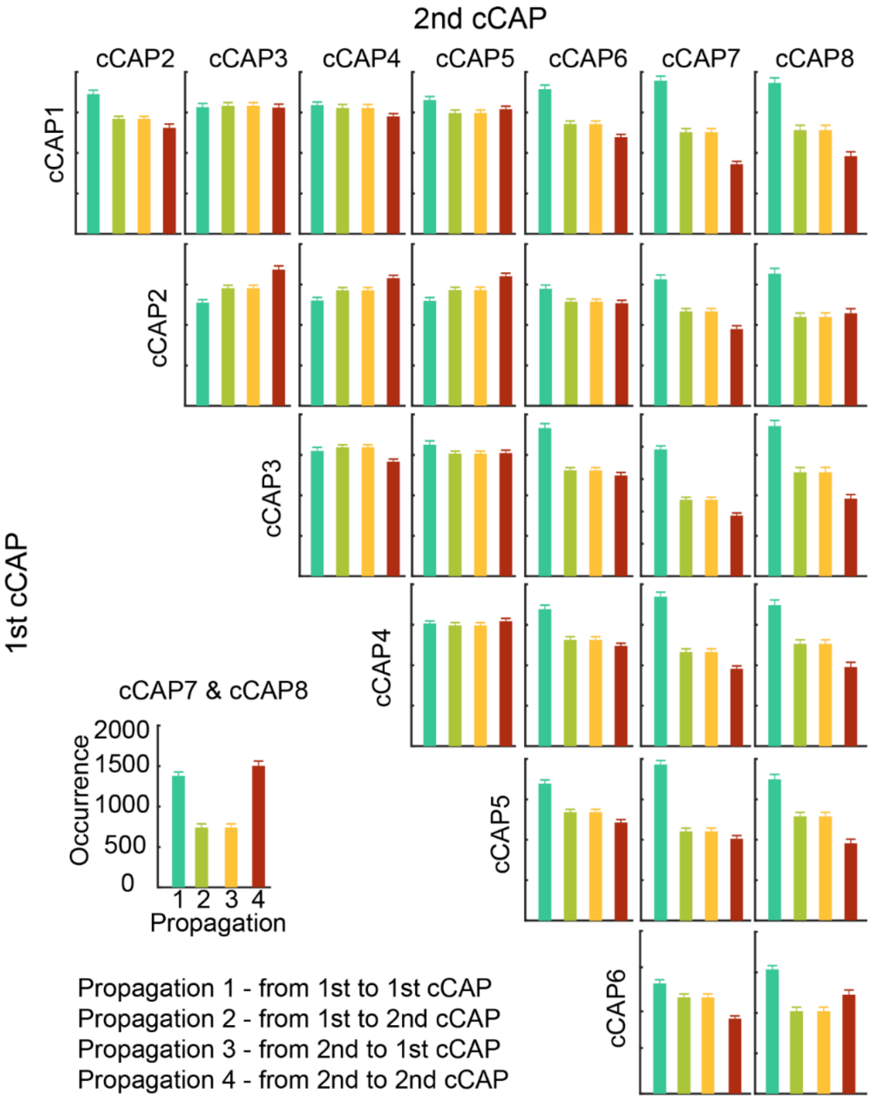


Supplementary Figure 7. Occurrences of the four types of propagations based on two *pseudo*-polarized states simulated by all possible pairs (total 27 pairs) of the eight cCAPs other than the CAPs 7-8 pair. Each panel represents one such pair (the 1$^{st}$ cCAP is in the y-axis and the 2$^{nd}$ cCAP is in the x-axis). For the sake of comparisons, the results based on two real polarized states (cCAPs 7-8) are also listed at the left bottom corner. It is noted that the phenomenon of the higher occurrences of two propagations between two polarized states as compared to the occurrences of two contrasting propagations is not observed in these simulated pairs of *pseudo*-polarized states. And the phenomena of the similar level for the occurrences of two contrasting propagations are also not observed in most simulated pairs of *pseudo*-polarized states.

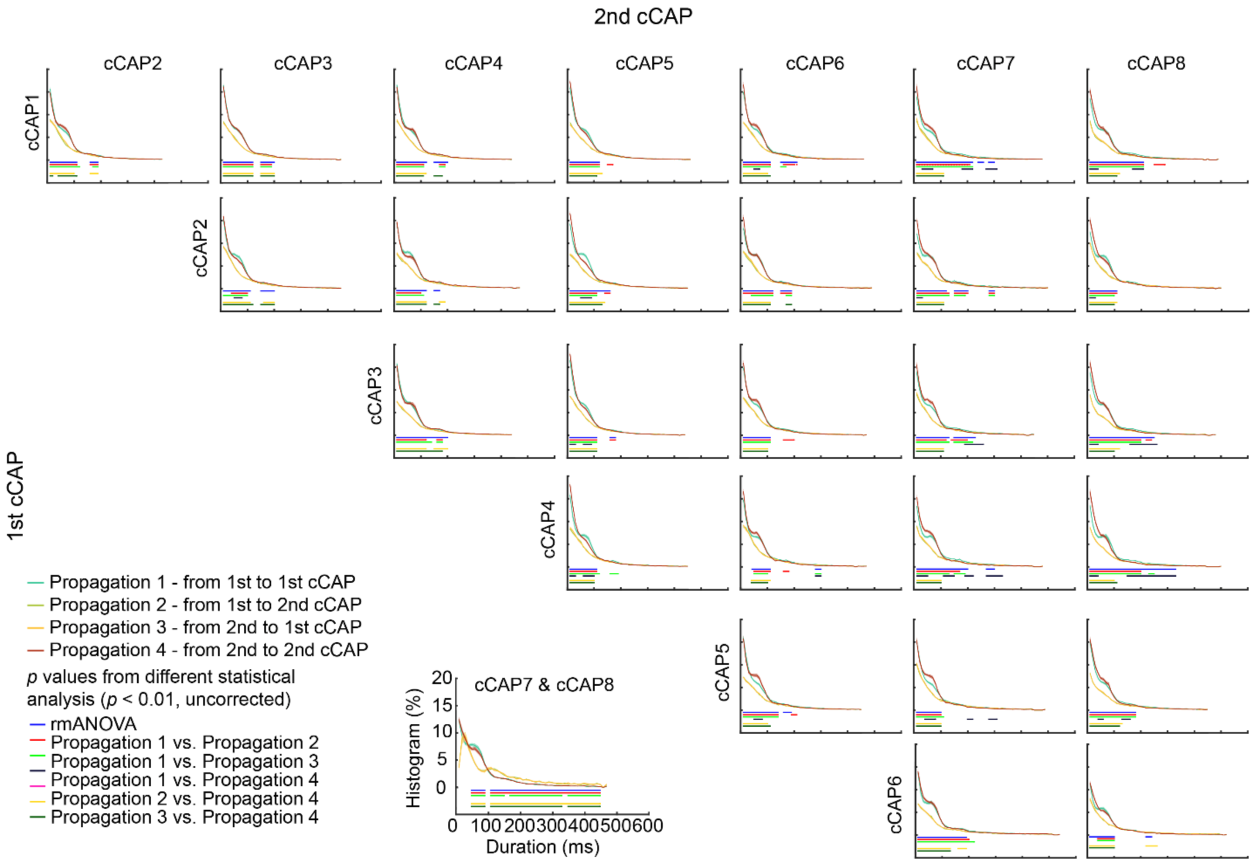


Supplementary Figure 8. Histograms of durations (SEM: shaded areas) of the four types of propagations as functions of duration time for the propagations based on two *pseudo*-polarized states simulated by all possible pairs (total 27 pairs) of the eight cCAPs other than the CAPs 7-8 pair. Each panel represents one such pair (the 1$^{st}$ cCAP is in the y-axis and the 2$^{nd}$ cCAP is in the x-axis). For the sake of comparisons, the result based on two real polarized states (cCAPs 7-8) is also listed at the left bottom corner. It is noted that there are no observations of characteristic peaks around 30ms and 100ms, as well as elevated occurrences of the propagations beyond 100ms between the simulated pairs of *pseudo*-polarized states, which are evident in the CAPs 7-8 pair as the real polarized states.

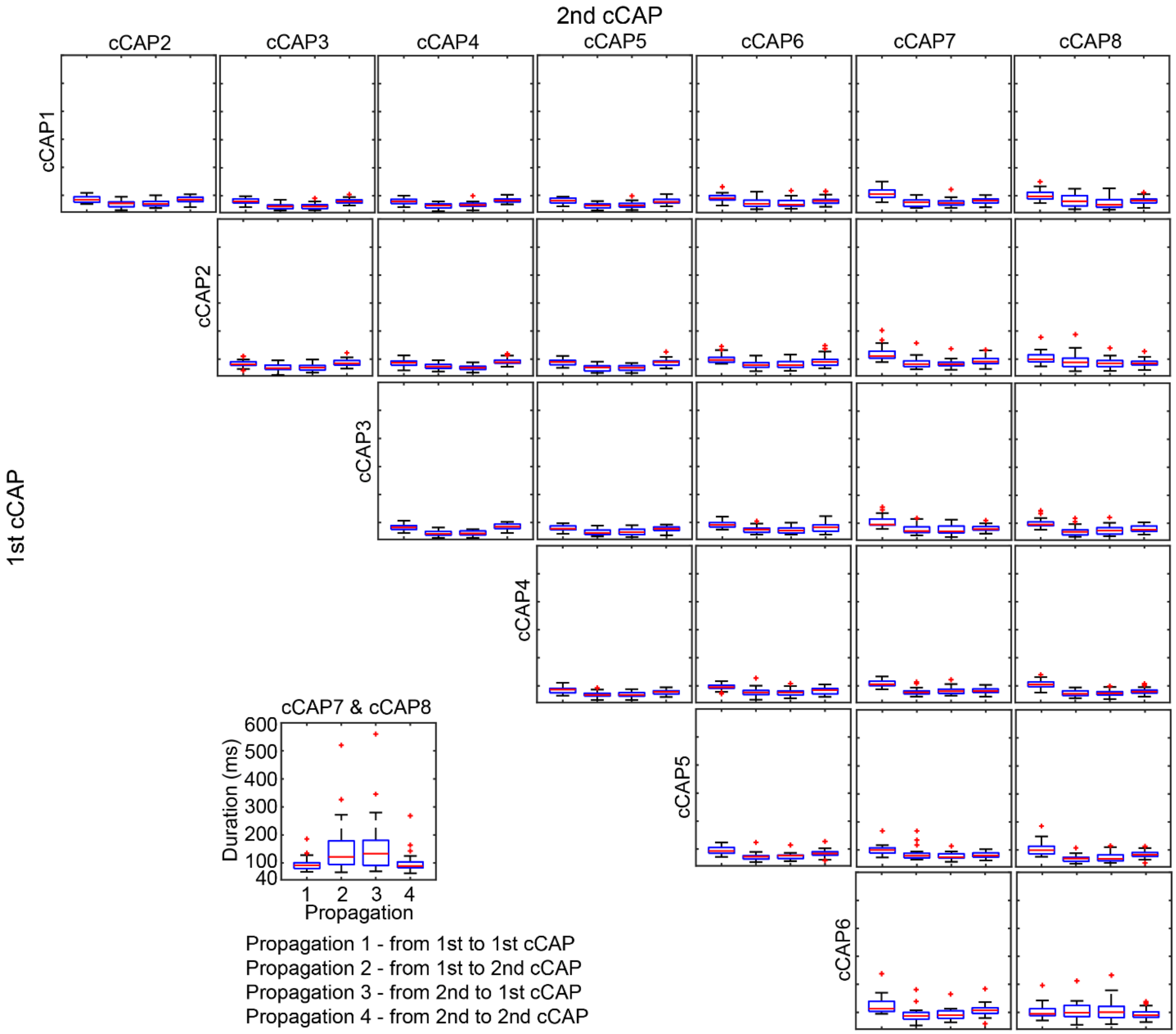


Supplementary Figure 9. Boxplots of durations for the four types of propagations based on two *pseudo*-polarized states simulated by all possible pairs (total 27 pairs) of the eight cCAPs other than the CAPs 7-8 pair. Each panel represents one such as pair (the 1$^{st}$ cCAP is in the y-axis and the 2$^{nd}$ cCAP is in the x-axis). For the sake of comparisons, the result based on two real polarized states (cCAPs 7-8) is also listed at the left bottom corner. It is noted that the durations of the two contrasting propagations are observed usually longer than the durations of the propagations between the simulated pairs of *pseudo*-polarized states, which is opposite to the observations in the cCAPs 7-8 pair as the real polarized states.

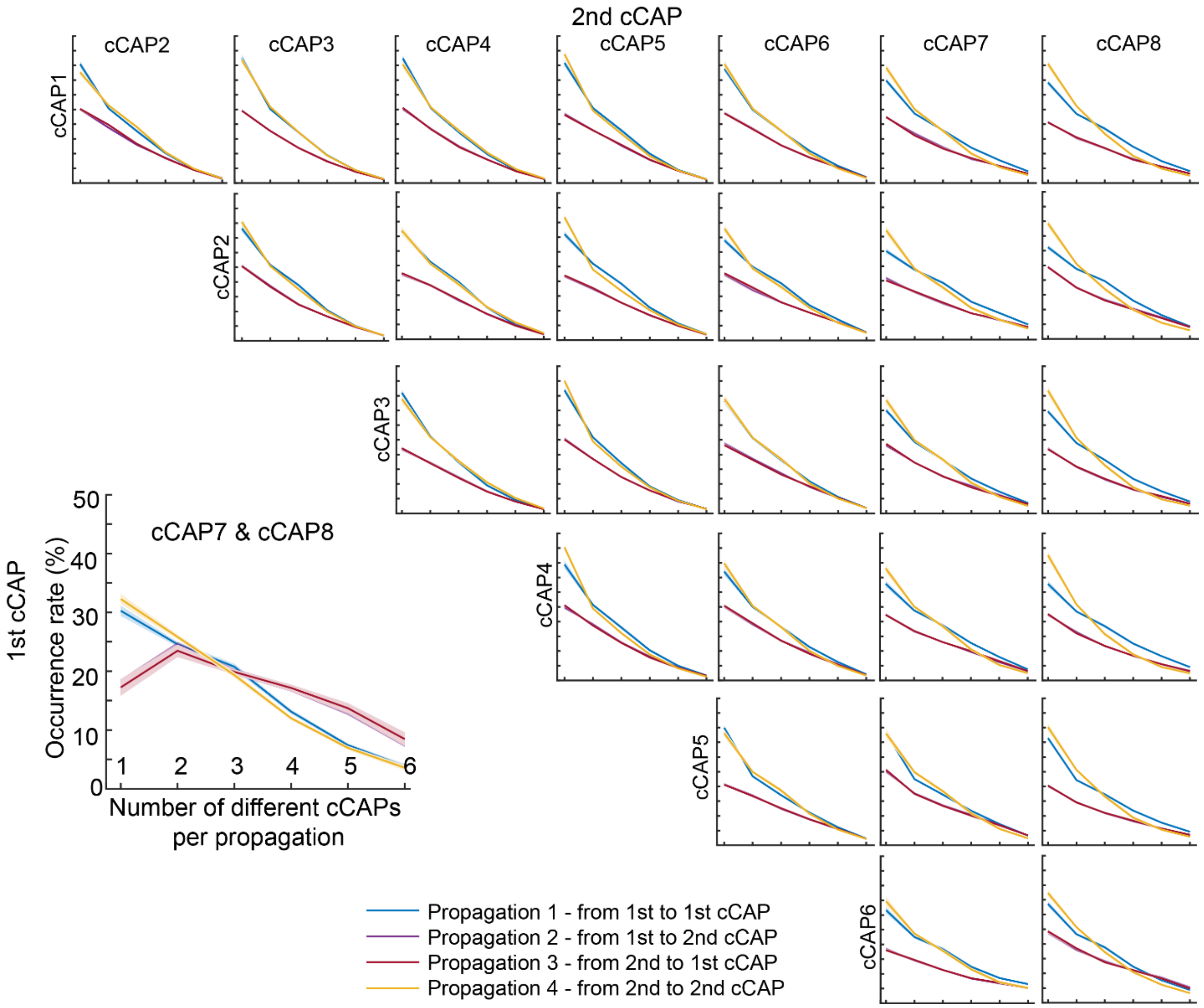


Supplementary Figure 10. Occurrence rates (SEM: shaded areas) of the four types of propagations as functions of the number of different cCAPs visited (other than two *pseudo*-polarized states) based on two *pseudo*-polarized states simulated by all possible pairs (total 27 pairs) of the eight cCAPs other than the CAPs 7-8 pair. Each panel represents one such as pair (the 1$^{st}$ cCAP is in the y-axis and the 2$^{nd}$ cCAP is in the x-axis). For the sake of comparisons, the result based on two real polarized states (cCAPs 7-8) is also listed at the left bottom corner. It is noted that elevated occurrence rates for the propagations between two *pseudo*-polarized states as compared with the two contrasting propagations for the number of different cCAPs visited as 4, 5, and 6 are absent, which are evident of statistical significance in the cCAPs 7-8 pair as the real polarized states.

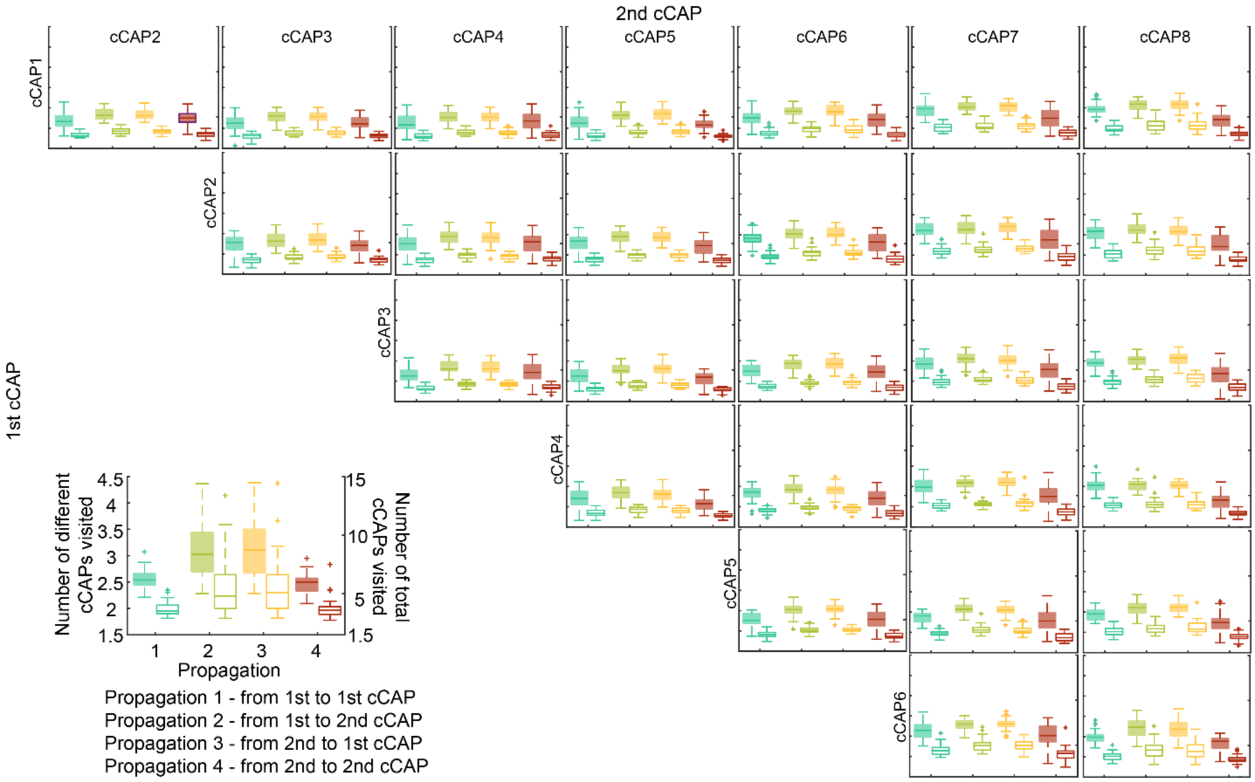


Supplementary Figure 11. Participant-level means (±SEM) of numbers of different cCAPs (left y-axis and boxplots with solid-fills) and total numbers of cCAPs (right y-axis and boxplots with no-fills) visited per propagation, based on two *pseudo*-polarized states simulated by all possible pairs (total 27 pairs) of the eight cCAPs other than the CAPs 7-8 pair. Each panel represents one such as pair (the 1$^{st}$ cCAP is in the y-axis and the 2$^{nd}$ cCAP is in the x-axis). For the sake of comparisons, the result based on two real polarized states (cCAPs 7-8) is also listed at the left bottom corner. It is noted that no similar results are observed as those observed with the cCAPs 7-8 pair as the real polarized states.

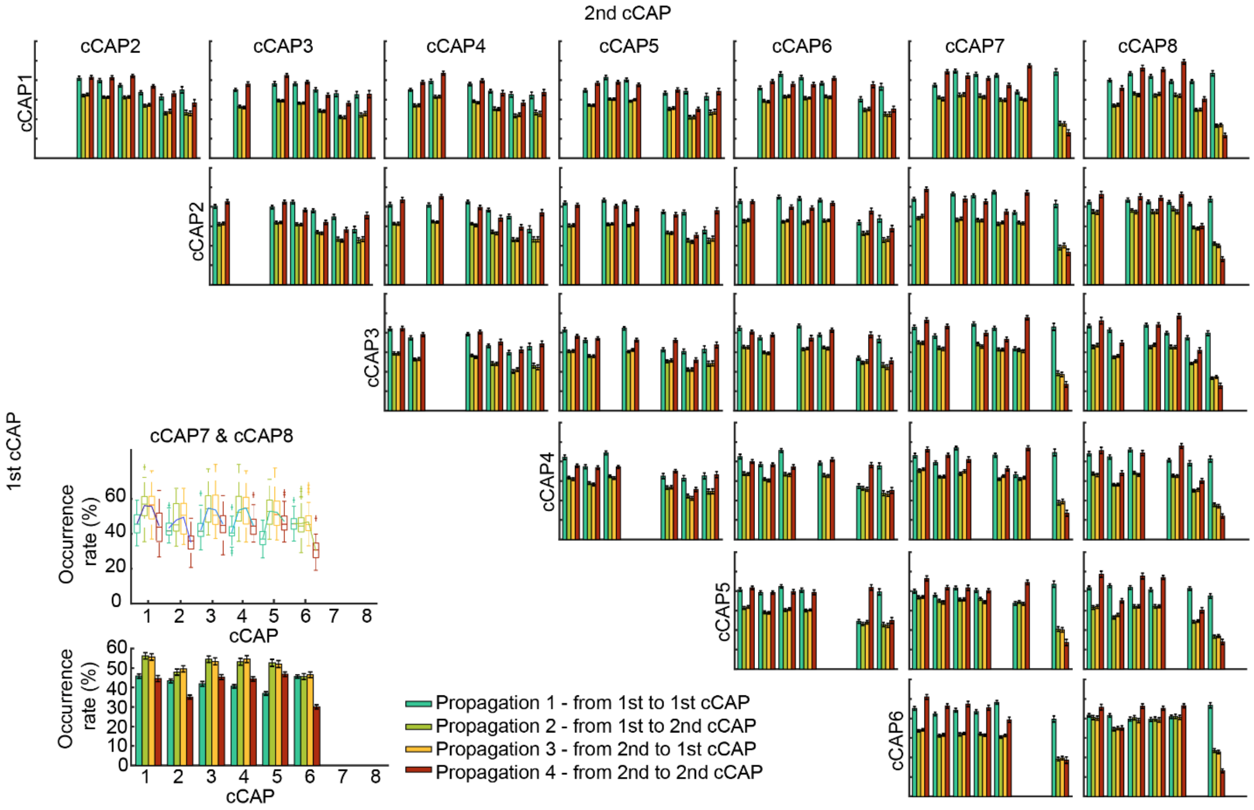


Supplementary Figure 12. Occurrence rates of the other six cCAPs within each type of propagation, based on two *pseudo*-polarized states simulated by all possible pairs (total 27 pairs) of the eight cCAPs other than the CAPs 7-8 pair. Each panel represents one such as pair (the 1$^{st}$ cCAP is in the y-axis and the 2$^{nd}$ cCAP is in the x-axis). For the sake of comparisons, the result based on two real polarized states (cCAPs 7-8) is also listed at the left bottom corner. It is noted that, with the cCAPs 7-8 pair as the polarized states, all other six cCAPs occur more in the two propagations between the two polarized states than in the two contrasting propagations. Meanwhile, with a pair of *pseudo*-polarized states, the rest six cCAPs usually occur more in the two contrasting propagations than in the two propagations between the two *pseudo*-polarized states.
